# Morph-specific patterns of sex organ positions in species with style length polymorphism

**DOI:** 10.1101/2021.04.25.441381

**Authors:** Shatarupa Ganguly, Deepak Barua

## Abstract

In style length polymorphism, morph-specific sequence of encountering male and female sex organs within a flower by pollinators can cause differences in the need to avoid self-pollination and encourage inter-morph pollination. We asked if this difference can lead to disparity in stigma-anther separation (herkogamy) between morphs and spatial match between sex organs of complementary morphs (reciprocity). Further, we tested if herkogamy, and hence the level of selfing, is fairly constant among individuals of a population. Additionally, we examined the relationship between herkogamy and reciprocity among individuals of a population to understand functional interactions between these two morphological traits. Using data on sex organ heights for >200 heterostylous species from the literature, we observed that the short-styled morph had higher herkogamy as compared to the long-styled morph indicating a higher need to avoid selfing. Reciprocity did not show a consistent difference between the upper and lower sex organs implying a strong influence of local ecological factors. In most populations, allometric relationships suggested that herkogamy and hence the level of selfing remains constant. Finally, we observed that herkogamy and reciprocity can be related among individuals of a population, sometimes indicating a potential trade-off between avoidance of self-pollination and facilitation of inter-morph pollination.

## INTRODUCTION

Sex organ positions in heterostylous plants are considered to play an important role in efficient inter-morph pollen transfer and hence maintenance of this style length polymorphism (Barrett & Shore, 2008). The differences in the relative position of anther and stigma between morphs causes differences in their physical interaction to various pollinators giving rise to morph-specific pollen pick up and deposition patterns. Morph-specific patterns in pollen transfer can result in differences in self, intra-morph and inter-morph pollen exchange, and subsequently influence selection for higher or lower herkogamy in morphs or reciprocity between complementary sex organs. General patterns of relative sex organ positions within and between morphs can potentially reveal the morphological basis of predominant morph-specific strategies of intra- and inter-morph pollen exchange. However, despite the presence of a large amount of information on sex organ positions in the extant literature, predictions related to general patterns of differences in herkogamy and reciprocity over a large number of heterostylous taxa have not been tested.

Heterostyly and stigma-height dimorphism are kinds of style-length polymorphism which is defined as the presence of two or three distinct floral morphs within a population differing in style lengths (Barrett, Jesson, & Baker, 2000). In heterostyly, the morphs additionally differ in anther positions resulting in reciprocal arrangement of sex organs between complementary morphs, a feature stigma-height dimorphism lacks (Baker, 2000; Barrett, 2002). In distyly (dimorphic heterostyly) both morphs exhibit ordered herkogamy where spatially separated male and female sex organs are encountered in a specific sequence. In the long-styled morph, the pollinator encounters the stigma first and then the anthers akin to approach herkogamy where the stigma is higher than the anthers in a flower (Webb & Lloyd, 1986; Opedal, 2018). In the short-styled morph the anthers are presented before the stigma to the pollinators similar to reverse herkogamy where the anther is higher than the stigma in a flower. Therefore, there is a higher chance of self-pollination in the short-styled morph as the pollinator can pick up pollen from the anthers before reaching the stigma of the same flower (Webb & Lloyd, 1986). This can lead to pollen discounting, stigma clogging, and in the absence of absolute heteromorphic incompatibility can also result in illegitimate fertilization (Zhou *et al*., 2015). Consequently, there can be a potential reduction, especially in female fitness, of the short-styled morph. Hence, the short-styled morph is likely to have higher herkogamy than the long-styled morph to compensate for the higher chance of self-pollination.

As most plants with stye length polymorphism have narrow corolla tubes, long-tongued pollinators are essential to ensure pollen transfer between the upper-level long-styled stigma and the short-styled anther as well as the lower level short-styled stigma and the long-styled anther (Ganders, 1979; Lloyd & Webb, 1992a; Simón-Porcar, Santos-Gally, & Arroyo, 2014). The upper-level sex organs can come in contact with most floral visitors as they are situated at the collar of, or exserted from the corolla tube (Nishihiro *et al*., 2000). Hence pollen exchange between the anther and stigma of the upper level can take place more frequently than the lower level. In fact, the long-styled stigma commonly exhibits higher total pollen loads than the short-styled stigma tube (Nicholls, 1985; Stone & Thomson, 1994; Liu, Wu, & Huang, 2016; Jacquemyn, Gielen, & Brys, 2018). The deposition of surplus pollen on the long-styled stigma might result in relaxed selection for high reciprocity between the sex organs of complementary morphs of the upper level (Haddadchi, 2013).

Fundamental differences in relative sex organ positions between morphs can also bring about morph-specific patterns of intrapopulation variation in herkogamy and consequently pollen flow. Herkogamy is a composite trait which is derived from the male and female sex organ positions within a flower (Opedal *et al*., 2017). Allometric changes in stigma and anther heights with respect to the corolla tube have been used to quantify the relative contribution of male and female sex organs towards changes in herkogamy (Richards & Koptur, 1993; Faivre & McDade, 2001). To maintain the same degree of herkogamy, changes in anther and stigma positions with changes in flower size within a population are expected to be isometric. However, if the change in size any of the sex organs is accelerated or deaccelerated relative to the other sex organ, herkogamy will vary across individuals. Since larger organs are expected to show higher variance (Runions & Geber, 2000; Opedal *et al*., 2017), the long-styled stigma and the short-styled anthers can exhibit greater changes in size across individuals as compared to the short-styled stigma and the long-styled anther. However, variation in anther height can be restricted as the anthers in most heterostylous species are attached to the corolla tube (Ganders, 1979). Thus a change in anther position will have concurrent functional consequences related to a change in corolla tube dimensions (Faivre, 2000). Hence, both anther and stigma positions can influence variation in herkogamy in the short-styled morph. The relative contribution of the male and female sex organs to the intrapopulation variation in herkogamy will determine whether pollen export or pollen receipt is more affected as a consequence of its impact on how sex organs come in contact with the pollinator’s body (Herlihy & Eckert, 2007).

The extent of intra- and inter-morph pollen exchange is determined by both herkogamy and reciprocity (Keller, Thomson, & Conti, 2014). Differences in herkogamy among individuals of a population can be related or unrelated to differences in reciprocity of the same individuals with sex organs of the complementary morph. Such a relationship can reveal how a change in organ heights that results in reduced or increased herkogamy and associated changes in the chance of self-pollination will simultaneously influence reciprocity and the chance of inter-morph pollination. Moreover, the relationship between herkogamy and reciprocity can also depend on the morph identity revealing selection pressures unique to a morph. The presence or lack of a relationship between herkogamy and reciprocity also has implications for understanding selection for reproductive assurance or against inbreeding depression.

In this study, we conducted a literature survey and collected data on sex organ position in dimorphic style length polymorphisms - distyly and stigma-height dimorphism. We performed phylogenetically corrected statistical analysis to ask whether mean herkogamy is higher in the short-styled morph indicating a greater need to avoid self-pollination. Similarly, we tested if mean reciprocity is lower in the upper sex organ level as compared to the lower sex organ level due to high pollen deposition on the long-styled stigma. To understand the potential morph-specific implications of intra-population variation in herkogamy on pollen flow, we asked if anther and stigma positions within flowers varied isometrically with respect to each other among individuals of a population. As described earlier, we expect the stigma position in the long-styled morph and both the anther as well as stigma position in the short-styled morph to have relatively greater contribution towards intra-population variation in herkogamy. Finally, we examined the relationship between herkogamy and reciprocity among individuals of populations to identify potential trade-offs between avoidance of self-pollination and inter-morph pollination.

## MATERIALS AND METHODS

### Literature Survey and data extraction

We conducted an exhaustive literature search using the keywords style length polymorphism, heterostyly, distyly and stigma-height dimorphism. To begin with, searches were performed using the ISI Web of Science and Google Scholar. Subsequently, we also conducted searches for the references cited in and cited by the initially identified publications. This study was restricted to the dimorphic character states of style length polymorphism, i.e., distyly and stigma-height dimorphism. The following data were extracted from these publications: (a) mean sex organ heights of both morphs for a population when reported for multiple individuals of each morph; (b) mean herkogamy when reported for multiple individuals for each morph; and, (c) anther and stigma heights of individuals of a population when this information was available for at least ten individuals of each morph. We extracted data from text, tables, scaled floral illustrations, and when necessary, by digitising graphs and figures using the software PlotDigitizer (http://plotdigitizer.sourceforge.net). The mean heights of the four sex organs were used to standardize herkogamy and reciprocity values to enable comparison across species with varying flower sizes. The mean species-level herkogamy data was used to make comparison between morphs. The data for anther and stigma heights of individuals were used to derive reciprocity and occasionally herkogamy when the mean herkogamy information was reported for a species.

### Calculation of Mismatch and herkogamy

As per convention (Lloyd & Webb, 1992b; Olesen *et al*., 2003), the long-styled stigma and the short-styled anther were defined as the upper level, and the short-styled stigma and the long-styled anther were defined as the lower level complementary sex organ positions. Reciprocity was calculated using the sex organ heights of individuals of a population as the mismatch between anther and stigma heights of the complementary morphs (Ganguly & Barua, 2020). To incorporate the effects of intra-population variation in sex organ heights, the absolute mismatch for every complementary stigma-anther pair was calculated and averaged for the upper and lower levels of sex organ positions. When mean herkogamy for both morphs of a population was not reported, it was calculated as the absolute difference in anther and stigma heights within a flower and averaged over individuals for each morph separately (Opedal, 2018). To understand which sex organ was relatively more important in bringing about morph-specific difference in herkogamy, we compared mean anther and stigma heights at each sex organ level. Paired t-test was performed to compare mean mismatch between sex organ levels, mean herkogamy between morphs, and mean anther and stigma heights at each sex organ level across species.

To understand how many species have higher herkogamy in the short-styled morph and higher mismatch in the upper-level sex organ as per our predictions, we performed independent t-test for each species to compare herkogamy and mismatch between morphs and levels, respectively. To this end, we derived herkogamy as the difference between stigma and anther heights of an individual for the species for which sex organ heights of individuals of a population were available. We calculated the mismatch of every individual as the average of the mismatch of the stigma of that individual with anthers of all the individuals of the complementary morph.

### Phylogenetic correction

We performed phylogenetically corrected paired t-test (Lindenfors, Revell, & Nunn, 2010) to examine differences in mismatch, herkogamy and sex organ heights between levels and morphs. Phylogenetically corrected statistical tests were performed using the R packages phytools (Revell, 2012) and ape (Paradis, Claude, & Strimmer, 2004). For the statistical analyses, species names were obtained from The Plant List (Version 1.1. Published on the Internet; http://www.theplantlist.org/, accessed on December 2018). The recent comprehensive phylogenetic tree for seed plants provided by Smith and Brown (2018) was used for the phylogenetically corrected statistical tests. Species which were not present in the tree provided by Smith and Brown (2018) were excluded from the analyses. When data for multiple populations of the same species were present, the population with the largest sample size was chosen for the analyses. Moreover, as differences in flower size among species could influence the results, mean herkogamy and mismatch for both morphs of each species were standardised using the grand mean of all four sex organ heights for that species. To assess the representation of different flower sizes in the data, we calculated the mean of average sex organ height of the upper level as a surrogate for flower size.

### Allometric relationships

In this section we used data on sex organ heights of individuals of a population when sex organ heights for at least 30 individuals of each morph were available. Some species were represented by multiple populations as long as the criterion of sample size was met. We examined the allometric relationships between anther and stigma heights of individuals of a population. The allometry between anther and stigma heights was examined by quantifying the slope of ranged major axis (RMA) type II regression using lmodel2 package (Legendre, 2018) of R (*ver 4*.*0*.*0*) (R Core Team, 2020). RMA regressions were used because of different variances in the estimates of anther and stigma heights (Legendre, 1998). We used *p-values* and 95% CI of the slope estimate to ask whether the slope was significantly different from one to test if the anther and stigma heights were isometrically related to each other.

### Relationship between herkogamy and reciprocity

The populations used to study allometric relationships between anther and stigma heights were also used to understand the relationship between herkogamy and reciprocity. We calculated mismatch for an individual as the average of all the mismatches between stigma height of that individual and anther heights of all individuals of the complementary morph. This measure of reciprocity reflects the chances of pollen deposition on the stigma of that individual and hence is a measure of female fitness. To examine the relationship between herkogamy and reciprocity, we calculated Pearson’s correlation coefficient between herkogamy and mismatch among individuals of each morph of a population.

## RESULTS

### Details of the extracted data

The final number of references used for this study after applying the selection criteria was 87. The initial data for mean herkogamy included 249 distylous species represented by 459 populations, and 23 species with stigma height dimorphism represented by 44 populations. Information for anther and stigma heights of individuals of a population of a species was available for 119 distylous species (211 populations) from 14 families, and 17 species (34 populations) from 7 families with stigma-height dimorphism. Overall, the number of species with distyly and stigma-height dimorphism used in this study represented 18 of 28 families (Naiki, 2012) for which heterostyly has been reported till date (Supplementary information Fig. S1). The range of flower sizes examined in this study (Supplementary information Fig. S1) was representative of the range of flower sizes reported for heterostylous species (Ganders, 1979).

### Difference in herkogamy between morphs

Out of the total number of species for which data for mean herkogamy for the long- and short-styled morphs was available, 230 distylous species and all 18 species with stigma-height dimorphism were present in the phylogenetic tree of seed plants provided in Smith & Brown (2018) (Supplementary information Table S1). Only these species were used for the phylogenetic corrected analyses. High variation in herkogamy was observed for both morphs especially in distylous species ranging from 0.39% to 155.10% of the grand mean of sex organ heights of both morphs. As expected, mean herkogamy was significantly higher in the short-styled morph compared to the long-styled morph in distylous species (*t* = 3.12, *df* = 229, *p*-value = 0.002; Fig. 1). The outcome was consistent after phylogenetically corrected paired t-test was performed (*t* = 3.13, *df* = 227, *p*-*value* = 0.002; Fig. 1). Similarly, in species with stigma-height dimorphism, mean herkogamy was higher in the short-styled morph (*t* = 2.65, *df* = 17, *p-value* = 0.02; Fig. 1). However, phylogenetically corrected paired t-test revealed no significant differences in herkogamy for the two morphs (*t* = 0.17, *df* = 15, *p-value* = 0.87).

**Figure 1:**
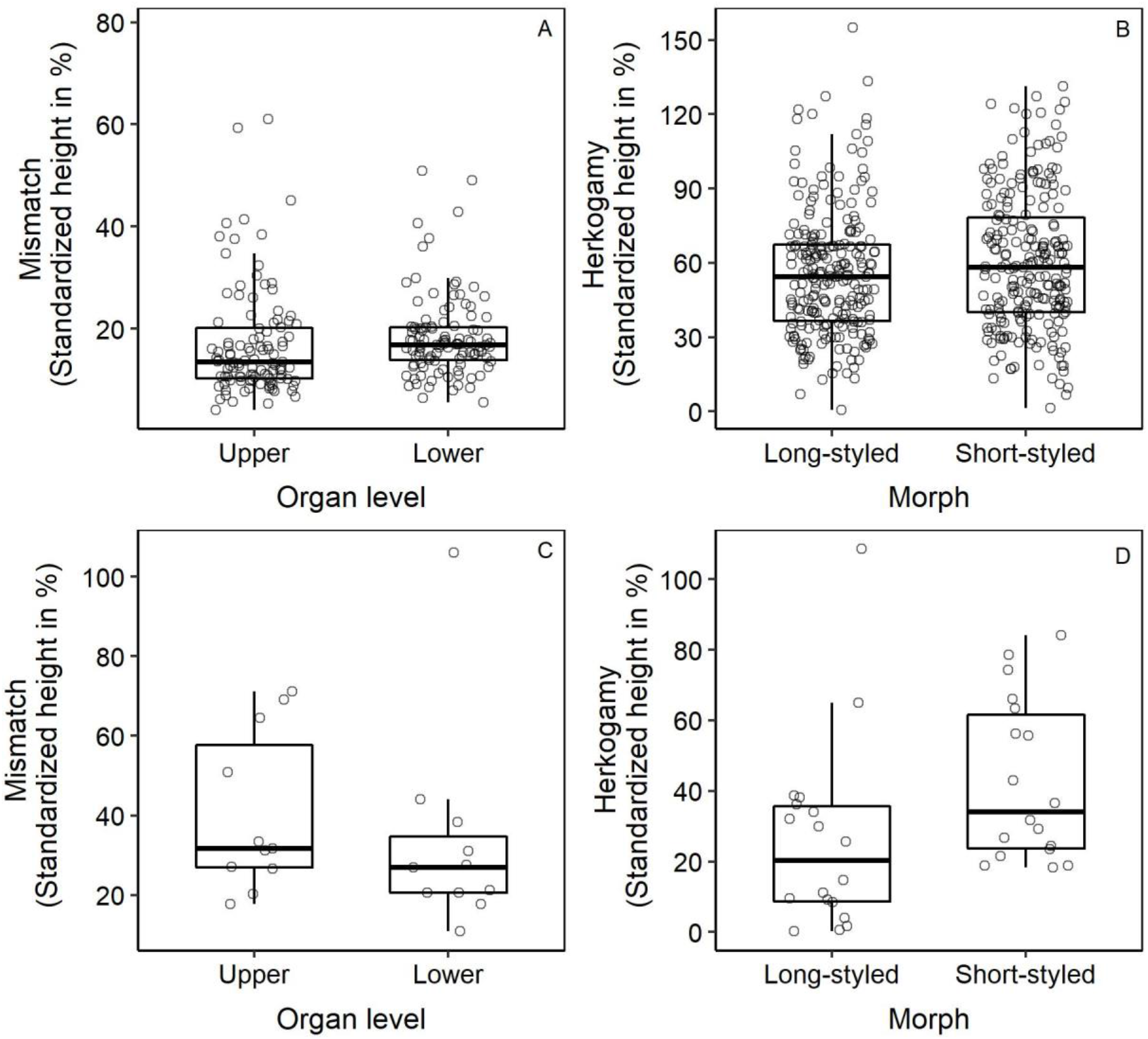
Mismatch (A and C) and herkogamy (B and D) in species with distyly (A and B) and stigma height dimorphism (C and D) presented as a percentage of the grand mean of all four mean sex organs heights. The sample sizes for herkogamy are 230 species for distyly and 18 species for stigma-height dimorphism. The sample sizes for mean mismatch are 107 species for distyly and 11 for stigma height dimorphism. The bold line in the boxplots represents the median, the ends of the box represent the first and the third quartiles and the whiskers represent 1.5 times the interquartile range.

Out of the 107 distylous species for which anther and stigma heights were available for individuals of the population, 53 species exhibited significantly higher mean herkogamy in the short-styled morph as compared to 35 species where the long-styled morph had higher herkogamy. In a total of 11 species with stigma-height dimorphism, 8 species displayed higher herkogamy in the short-styled morph as compared to 3 species which exhibited higher herkogamy in the long-styled morph.

Mean stigma height of the long-styled morph was marginally higher than mean anther height of the short-styled morph in distylous species (*t* = 1.89, *df* = 229, *p-value* = 0.06) but the difference was not significant on performing phylogenetically corrected paired t-test (*t* = 0.67, *df* = 227, *p-value* = 0.5). However, mean anther height of the long-styled morph was significantly higher than mean stigma height of the short-styled morph in distylous species when analysed without (*t* = 7.05, *df* = 229, *p-value* < 0.001) or with phylogenetic correction (*t* = 2.66, *df* = 227, *p-value* = 0.008). In species with stigma-height dimorphism, mean stigma height of the long-styled morph was not significantly higher than mean anther height of the short-styled morph (*t* = 1.59, *df* = 17, *p-value* = 0.13), while mean stigma height of the short-styled morph was significantly smaller than the mean anther height of the long-styled morph (*t* = -10.40, *df* = 17, *p-value* < 0.001). However, both were not statistically significant when phylogenetically corrected paired t -test was performed (*t* = 0.05, *df* = 15, *p-value* = 0.96 and *t* = -0.20, *df* = 15, *p-value* = 0.85, respectively).

### Difference in mean mismatch between sex organ levels

The final number of species which were present in the phylogenetic tree (Smith & Brown, 2018) and hence were used for statistical analyses was 107 for distylous species and 11 for species with stigma-height dimorphism. Mean mismatch between the upper and lower sex organ levels was not significantly different in species with distyly (*t* = 1.17, *df* = 106, *p-value* = 0.25; Fig. 1) or stigma height dimorphism (*t* = -0.73, *df* = 10, *p-value* = 0.48). These differences were also not significant after phylogenetically corrected paired t-test was performed for species with distyly (*t* = 1.19, *df* = 104, *p-value* = 0.24) as well as stigma-height dimorphism (*t* = -0.05, *df* = 8, *p-value* = 0.98).

Out of the abovementioned 107 distylous species, 63 species exhibited significantly higher mismatch in the upper level as compared to 30 species where the lower level had higher mismatch. In the 11 species with stigma-height dimorphism, 3 species displayed higher mismatch in the upper level as compared to 8 species which exhibited higher mismatch in the lower level.

### Allometric relationship between anther and stigma heights

The average slope of the relationship between stigma and anther heights of individuals was 1.10 ± 1.47 (mean ± 95% CI; *n* = 73 populations) for the long-styled morph, and 1.10 ± 0.27 (mean ± 95% CI; *n* = 79 populations) for the short-styled morph in distylous species. In species with stigma-height dimorphism, the average slope was 1.33 ± 0.48 (mean ± 95% CI; *n* = 17 populations) and 1.03 ± 0.73 (mean ± 95% CI; *n* = 14 populations) for the long- and short-styled morphs, respectively.

Out of the 50 long-styled and 55 short-styled distylous populations which exhibited a significant slope, 31 long-, and 39 short-styled populations had slopes which were not significantly different from one according to the 95% CI (Table 1 A). Higher change in stigma height as compared to the anther height was observed in 17 long-styled and 10 short-styled populations. On the other hand, greater increase in anther height as compared to stigma height was observed in one long-styled and four short-styled distylous populations. One long- and two short-styled populations negative relationship between anther and stigma heights.

**Table 1:** Details of the slope of the allometric relationship between stigma and anther heights of species with: A) distyly; and B) stigma-height dimorphism. Numbers denote the number of populations with RMA regression slope in that category according to the 95% CI. Each column represents the number of populations with that slope value with *P* < 0.05 for RMA regression. LS refers to the long-styled morph while SS refers to the short-styled morph.

|  | Morp<br>h | slope =<br>1 | slope <<br>0 | 0 < slope <<br>1 | slope ><br>1 |
| --- | --- | --- | --- | --- | --- |
| **A) Distyly** | LS | 31 | 1 | 17 | 1 |
|  | SS | 39 | 2 | 10 | 4 |
| **B) Stigma-height**<br>**dimorphism** | LS | 11 | 0 | 2 | 1 |
|  | SS | 7 | 0 | 1 | 1 |

Of the 15 long-styled and 9 short-styled populations with stigma-height dimorphism which exhibited a significant slope, 11 long-styled and 7 short-styled populations had slopes which were not significantly different from one according to the 95% CI (Table 1 B). A higher change in stigma height as compared to anther height was observed for two long-styled and one short-styled populations. A higher change in anther height as compared to stigma height was exhibited by one population of long- and short-styled morphs each. There were no populations which exhibited a negative relationship between anther and stigma heights. Overall, interestingly, the value of slope differed among different populations of the same species.

### Relationship between herkogamy and mismatch

Both long- and short-styled morphs of the distylous populations exhibited positive as well as negative relationships between herkogamy and mismatch. There was no significant difference in the number of positive and negative relationship in the long-styled (*χ*^*2*^ test: *p-value* = 0.63, *n* = 73 populations) and short-styled (*χ*^*2*^ test: *p-value* = 0.09, *n* = 79 populations) morphs in distylous populations (Table 2 A). On the other hand, species with stigma-height dimorphism had a significantly higher number of populations with a positive relationship in both long-styled (*χ*^*2*^ test: *p-value* < 0.001, *n* = 17 populations) and short-styled (*χ*^*2*^ test: *p-value* < 0.001, *n* = 14 populations) populations (Table 2 B).

**Table 2:** Relationship between herkogamy and mismatch in A) distyly and B) stigma-height dimorphism for the long- and the short-styled morph. Negative and positive denote negative or positive Pearson’s correlation coefficient between the two traits. The numbers denote the number of populations with the direction of the relationship followed by the number of populations with correlation coefficients with *P* < 0.05 in brackets.

| Character state | Morph | Negative | Positive |
| --- | --- | --- | --- |
| **A) Distyly** | Long-styled | 34 (22) | 39 (30) |
|  | Short-styled | 32 (12) | 47 (22) |
| **B) Stigma-height dimorphism** | Long-styled | 0 (0) | 17 (15) |
|  | Short-styled | 2 (0) | 12 (10) |

## DISCUSSION

The results from this study revealed how morph-specific selection pressures and developmental constraints can shape variation in herkogamy and reciprocity in species with distyly and stigma-height dimorphism. Herkogamy was higher in the short-styled morph possibly reflecting the importance of avoiding self-pollination especially in this morph. Reciprocity was not higher in the lower sex organ level as expected and examination at the species level showed no consistent pattern. This implies that reciprocity may be dependent on local factors, for example, the kind and abundance of visiting pollinators. As expected, allometric relationship between anther and stigma heights revealed that in most cases herkogamy was constant among individuals of a population. The allometric relationship also revealed that when herkogamy varied among individuals, the variation was largely driven by the stigma position of both morphs in distylous species. This suggests that in most cases, intra-population variation in herkogamy will possibly affect pollen receipt more than pollen export in both morphs. Both negative and positive relationships between herkogamy and reciprocity was observed implying that the relationship is shaped by local factors specific to a morph or population.

Understanding differences in sex organ heights between morphs has been a central subject of various studies on heterostylous species (Thompson & Dommee, 2000; Kudoh *et al*., 2001; Kálmán *et al*., 2007). Differences in herkogamy between morphs have been reported for a lot of species with distyly and stigma-height dimorphism (Thompson & Dommee, 2000; Cesaro *et al*., 2004; Li *et al*., 2010; Liu *et al*., 2012; Faife-Cabrera, Ferrero, & Navarro, 2014; Novo *et al*., 2018; Barranco, Arroyo, & Santos-Gally, 2019). Such differences between morphs are associated with differences in inter-morph pollen transfer due to avoidance of self- and intra-morph pollination (Baena-Díaz *et al*., 2012; Keller *et al*., 2014). Higher herkogamy has been observed in the short-styled morph in single species studies in species with distyly as well as stigma-height dimorphism. However, numerous examples of higher herkogamy in the long-styled morph are also available as evident in the results from this analysis of a larger number of species (Opler, Baker, & Frankie, 1975; Barrett & Richards, 1990; Massinga, Johnson, & Harder, 2005; Jones, 2012; Hernández-Ramírez, 2012; Haddadchi, 2013; Meeus *et al*., 2013; Faife-Cabrera *et al*., 2014). An important factor which was beyond the scope of this study is the presence and degree of heteromorphic incompatibility which, along with herkogamy, can reduce chances of illegitimate mating (Pailler & Thompson, 1997). Mechanisms like the presence of incompatibility and dichogamy can compensate for the loss in female fitness due to lower herkogamy (Cesaro *et al*., 2004; Simon-Porcar *et al*., 2015). The lack of information for the non-binary nature of the degree of incompatibility and the distinction between self, intra-morph and inter-morph incompatibility makes it difficult to compare among species hence inhibiting its inclusion in a study like this. That the majority of species in this analysis exhibited higher herkogamy in the short-styled morph suggests the order of presentation of anther and stigma remains a hindrance to increasing outcrossing irrespective of other selection pressures.

The comparison of mean sex organ heights at each level revealed that the significantly lower height of the short-styled stigma as compared to the long-styled anther of the lower sex organ level can be instrumental in achieving a higher herkogamy in the short-styled morph. This is perhaps because most distylous species have anthers attached to the corolla tube and hence changes in anther height may be more constrained. It has been suggested that the short-styled stigma is under opposing selection pressures for maintaining herkogamy and facilitating easier contact with pollinators by increasing its height (Nishihiro *et al*., 2000). This study indicates that maintaining herkogamy could be more important for the short-style morph of most distylous species.

Considered to be one of the primary factors facilitating inter-morph pollen transfer, reciprocity has been at the centre of studies dealing with heterostylous species. As with herkogamy, many studies have compared the reciprocity among the upper and lower sex organ levels (Lau & Bosque, 2003; Keller *et al*., 2016; Jacquemyn *et al*., 2018). Although more species exhibited lower reciprocity in the upper sex organ level, the overall results from our analysis were contrary to our expectation of lower reciprocity in the upper sex organ level. As pollinator size, tongue-length, behaviour and abundance can play a significant role in determining sex organ positions as well as inter-morph pollen transfer, they are possibly responsible for the observed lack of a pattern (Pérez-Barrales, Arroyo, & Scott Armbruster, 2007; Pérez-Barrales & Arroyo, 2010; Santos-Gally *et al*., 2013; Simón-Porcar *et al*., 2014). In fact, a study suggests that plants coevolve with efficient long-tongued pollinators and exhibit higher reciprocity when they are abundant or are the primary pollinators (Ferrero *et al*., 2011).

Floral coevolution with various pollinators determines the level of floral phenotypic integration or floral trait correlations (Berg, 1960). Species with style length polymorphism, mostly characterised with tubular corolla, are expected to be pollinated by specialised long-tongued pollinators leading to a relatively high trait correlation. As expected, in most species stigma and anther heights were positively related to each other. Surprisingly there were a few species where anther and stigma height were negatively related. The lack of a relationship or a weak relationship observed in some of the populations can be indication of pollination by a less efficient pollinator (Pérez-Barrales *et al*., 2007; Perez-Barrales *et al*., 2014). Isometry was observed in both morphs in most populations with distyly and stigma-height dimorphism indicating maintenance of the degree of herkogamy and the level of outcrossing. Pollen are deposited on specific points of pollinator’s body from where it is picked up by the stigma. Intra-population variation in sex organ position reduces the efficiency of pollen exchange due to inconsistency in the position onto or from which pollen is deposited or picked up from the pollinator’s body (Armbruster *et al*., 2017). Pollen export and outcross siring success are relatively more affected when anther position fluctuates more, whereas, outcross pollen deposition is more affected if stigma position exhibits more fluctuation (Herlihy & Eckert, 2007). Overall, in both long- and short-styled morphs, stigma height contributed more to variation in herkogamy than anther height irrespective of sex organ level (Herlihy & Eckert, 2007; Jiménez-López *et al*., 2019), as expected. This also indicates that stigma position is more amenable to changes in size unlike the anthers which are constrained by their attachment to the corolla tube (Faivre & McDade, 2001). As a consequence, stigma height in the long-styled as well as short-styled morph can readily respond to selection for higher outcross pollen deposition mediated by changes in herkogamy. Additionally, it suggests that the changes in anther height of both morphs in response to selection for altered pollen export will be hindered. The consequences of this inference for maintaining high reciprocity and sufficient inter-morph pollen flow in the lower sex organ level ultimately affecting the maintenance of the polymorphism should be further investigated.

The variation in herkogamy was related to variation in reciprocity in most of the study species. When the relationship between herkogamy and reciprocity is positive, plants can increase their capability to avoid self-pollen deposition while also increasing their chances of legitimate pollen transfer. But, when this relationship is negative, there will be a trade-off between the two functions. The former can be adaptive when plant populations are not pollen limited, but there is high inbreeding depression (Ushimaru & Nakata, 2002). However, when populations are pollen limited and need reproductive assurance (Ashman *et al*., 2004), a negative relationship will be a better strategy as it will ensure higher reciprocity with lower herkogamy and hence maximum reproductive assurance provided that heteromorphic incompatibility is absent. Differences in the direction of the relationship between morphs of a population and among populations is perhaps a manifestation of differences in local ecological scenarios and how that influences each morph.

In this study, we put together information regarding sex organ positions for a large number of species with style length polymorphism which helped us understand general patterns in herkogamy and reciprocity at the level of populations as well as individuals. One of the most important conclusions of this study is that the difference in order of presentation of sex organs between morphs can significantly influence difference in traits like herkogamy which are known to influence levels of inbreeding and outcrossing. Our results reveal that developmental causes can determine the likely consequence of intra-population variation in herkogamy on pollen export or receipt. Interestingly, we show that herkogamy and reciprocity can be related and a trade-off between avoidance of self-pollination and promotion of inter-morph pollination can exist. Future work that includes information on pollinators, incompatibility, pollen load and reproductive output etc. would help consolidate the results from this study. Such studies are needed to comprehensively understand the adaptive significance of sex organ positions in the maintenance of these polymorphisms. The lack of clear results in species with stigma-height dimorphism is likely due to the low sample size, a reflection of the handful of studies conducted on these species. Understanding the functional significance of sex organ positions in species with stigma-height dimorphism is crucial to unravel the causes of evolution and maintenance of heterostyly and hence demands more attention.

## AUTHOR CONTRIBUTIONS

SG and DB conceived the idea and extracted the data. SG analysed the data. SG and DB wrote the manuscript.

