## Supplementary Table S1 for "Morph-specific patterns of sex organ positions in species with style length polymorphism"

### Supplementary information

**Table S1:** Mean and SD (in mm) of stigma height (Stigma), anther height (Anther), herkogamy (HER) and mismatch (MM) of the species used in this study. Style length polymorphisms (PM) included are distyly (D) and stigma-height dimorphism (SHD). L and S represent the long- and short-styled morphs (M), respectively for mean sex organ heights and mean herkogamy. UP and LO represent upper and lower sex organ level (SL) for mean mismatch. Sample size is denoted by *n*. Reference denotes the serial number for the study from which data was extracted as per References at the end of this table. Three distylous species (*Arcytophyllum macbridei*, *Polygonum hastatosagittatum and Psychotria trichophora*) which were not identified by the phylogenetic tree but were included in evaluation of allometric relationship between anther and stigma heights have also been included in this table.

| Species | PM | M/SL | *n* | Stigma | SD | Anther | SD | HER | SD | MM | SD | Ref |
|---|---|---|---|---|---|---|---|---|---|---|---|---|
| *Arcytophyllum aristatum* | D | L/UP | 12 | 5.40 | 0.50 | 3.80 | 0.20 | 1.70 | 0.50 | 4.66 | 0.36 | 1 |
| Standl. | | S/LO | 12 | 4.20 | 0.20 | 5.50 | 0.50 | 1.30 | 0.40 | 0.46 | 0.29 | |
| *Arcytophyllum capitatum* | D | L/UP | 17 | 7.50 | 1.00 | 3.40 | 0.50 | 4.10 | 1.20 | 1.27 | 1.13 | 1 |
| (Benth.) K.Schum. | | S/LO | 16 | 3.60 | 0.60 | 6.30 | 0.70 | 2.70 | 0.90 | 0.58 | 0.47 | |
| *Arcytophyllum ciliolatum* | D | L/UP | 12 | 4.60 | 0.50 | 2.60 | 0.30 | 2.00 | 0.40 | 0.70 | 0.50 | 1 |
| Standl. | | S/LO | 12 | 2.30 | 0.40 | 4.80 | 0.70 | 2.50 | 0.60 | 0.49 | 0.31 | |
| *Arcytophyllum filiforme* | D | L/UP | 10 | 5.30 | 1.00 | 4.00 | 0.20 | 1.30 | 0.60 | 1.23 | 0.83 | 1 |
| (Ruiz & Pav.) Standl. | | S/LO | 10 | 3.40 | 0.60 | 4.70 | 0.90 | 1.30 | 0.80 | 0.92 | 0.64 | |
| *Arcytophyllum lavarum* | D | L/UP | 104 | 5.00 | 0.80 | 3.50 | 0.70 | 2.70 | 0.21 | 0.62 | 0.50 | 1, 2 |
| K.Schum. | | S/LO | 100 | 2.70 | 0.50 | 5.60 | 0.80 | 1.98 | 0.22 | 0.64 | 0.50 | |
| *Arcytophyllum macbridei* | D | L/UP | 30 | 9.40 | 1.00 | 5.70 | 0.60 | 3.70 | 1.00 | 1.38 | 0.97 | 1 |
| Standl. | | S/LO | 23 | 5.20 | 0.40 | 8.20 | 0.70 | 3.00 | 0.70 | 0.78 | 0.57 | |
| *Arcytophyllum rivetii* | D | L/UP | 11 | 5.60 | 0.80 | 3.50 | 0.60 | 2.00 | 0.60 | 0.77 | 0.59 | 1 |
| Danguy & Cherm. | | S/LO | 10 | 3.00 | 0.40 | 5.30 | 0.60 | 2.20 | 0.60 | 0.59 | 0.59 | |
| *Arcytophyllum setosum* | D | L/UP | 10 | 8.10 | 0.80 | 5.40 | 0.60 | 2.80 | 0.80 | 1.13 | 0.76 | 1 |
| (Ruiz & Pav.) Schltdl. | | S/LO | 10 | 4.20 | 0.80 | 7.40 | 0.80 | 3.20 | 0.70 | 1.31 | 0.79 | |

Table 1 continued.

| **Species** | **PM** | **M/SL** | ***n*** | **Stigma** | **SD** | **Anther** | **SD** | **HER** | **SD** | **MM** | **SD** | **Ref** |
|---|---|---|---|---|---|---|---|---|---|---|---|---|
| *Arcytophyllum thymifolium* | D | L/UP | 11 | 6.70 | 0.60 | 3.10 | 0.10 | 3.50 | 0.50 | 0.81 | 0.57 | 1 |
| (Ruiz & Pav.) Standl. | | S/LO | 10 | 3.40 | 0.80 | 6.40 | 0.70 | 3.00 | 0.80 | 0.54 | 0.43 | |
| *Arcytophyllum vernicosum* | D | L/UP | 14 | 5.10 | 0.50 | 2.80 | 0.40 | 2.30 | 0.50 | 0.83 | 0.59 | 1 |
| Standl. | | S/LO | 14 | 3.20 | 0.40 | 5.00 | 0.90 | 1.80 | 1.10 | 0.56 | 0.43 | |
| *Bouvardia ternifolia* | D | L/UP | 25 | 27.50 | 2.60 | 20.70 | 2.50 | 6.60 | 1.00 | 3.05 | 2.33 | 3, 4 |
| (Cav.) Schltdl. | | S/LO | 25 | 21.00 | 2.10 | 28.40 | 2.70 | 7.40 | 2.30 | 2.58 | 1.96 | |
| *Carapichea ipecacuanha* | D | L/UP | 31 | 7.20 | 0.80 | 3.90 | 0.40 | 3.21 | 0.49 | 1.02 | 0.70 | 5 |
| (Brot.) L.Andersson | | S/LO | 33 | 3.20 | 0.50 | 6.40 | 0.70 | 3.11 | 0.55 | 0.68 | 0.46 | |
| *Chassalia corallioides* | D | L/UP | 25 | 13.18 | 1.23 | 10.79 | 1.39 | 3.45 | 1.06 | 2.33 | 1.67 | 6 |
| (Cordem.) Verdc. | | S/LO | 25 | 7.61 | 1.46 | 14.00 | 1.60 | 9.35 | 1.82 | 4.36 | 1.64 | |
| *Cordia dodecandra* | D | L/UP | 8 | 38.60 | 2.70 | 27.90 | 1.60 | 7.90 | 2.50 | - | - | 7 |
| A.DC. | | S/LO | 10 | 26.10 | 3.20 | 35.31 | 4.00 | 5.70 | 3.70 | - | - | |
| *Cordia macrocephala* | D | L/UP | 22 | 13.69 | 1.35 | 9.15 | 1.13 | 4.54 | 1.16 | 1.81 | 1.18 | 8 |
| (Desv.) Kunth | | S/LO | 17 | 11.27 | 0.94 | 12.37 | 1.12 | 1.12 | 0.71 | 2.14 | 1.40 | |
| *Cordia sebestena* | D | L/UP | 9 | 31.30 | 2.70 | 25.80 | 2.40 | 6.00 | 1.90 | - | - | 7 |
| L. | | S/LO | 11 | 25.30 | 2.20 | 31.90 | 2.10 | 5.30 | 2.40 | - | - | |
| *Coussarea platyphylla* | D | L/UP | 50 | 43.18 | 9.31 | 37.76 | 7.99 | 9.73 | 7.22 | 9.11 | 6.70 | 9 |
| Müll.Arg. | | S/LO | 47 | 28.31 | 3.42 | 45.24 | 6.29 | 17.03 | 5.92 | 11.01 | 6.50 | |
| *Damnacanthus indicus* | D | L/UP | 11 | 11.68 | 1.43 | 8.83 | 0.99 | 2.85 | 0.75 | 1.63 | 1.13 | 10 |
| C.F.Gaertn. | | S/LO | 11 | 7.50 | 0.89 | 10.63 | 1.04 | 3.13 | 0.75 | 1.55 | 0.98 | |

Table 1 continued.

| **Species** | **PM** | **M/SL** | ***n*** | **Stigma** | **SD** | **Anther** | **SD** | **HER** | **SD** | **MM** | **SD** | **Ref** |
|---|---|---|---|---|---|---|---|---|---|---|---|---|
| *Damnacanthus major* | D | L/UP | 10 | 13.48 | 1.65 | 7.57 | 1.20 | 5.92 | 1.71 | 2.58 | 1.65 | 10 |
| Siebold & Zucc. | | S/LO | 12 | 8.56 | 1.35 | 11.05 | 1.04 | 2.49 | 1.06 | 1.63 | 1.14 | |
| *Declieuxia fruticosa* | D | L/UP | 84 | 5.46 | 0.71 | 3.22 | 0.43 | 2.20 | 0.56 | 0.90 | 0.65 | 11 |
| (Willd. ex Roem. & Schult.) Kuntze | | S/LO | 90 | 3.53 | 0.59 | 5.53 | 0.81 | 2.01 | 0.79 | 1.05 | 0.57 | |
| *Dionysia aretioides* | D | L/UP | 2 | 14.50 | - | 7.75 | - | 6.75 | - | - | - | 12 |
| (Lehm.) Boiss. | | S/LO | 2 | 7.75 | - | 15.50 | - | 7.75 | - | - | - | |
| *Dionysia bryoides* | D | L/UP | 2 | 11.00 | - | 4.75 | - | 6.25 | - | - | - | 12 |
| Boiss. | | S/LO | 2 | 4.75 | - | 9.50 | - | 4.75 | - | - | - | |
| *Dionysia hissarica* | D | L/UP | 2 | 14.00 | - | 7.00 | - | 7.00 | - | - | - | 12 |
| Lipsky | | S/LO | 2 | 7.00 | - | 14.00 | - | 7.00 | - | - | - | |
| *Dionysia lurorum* | D | L/UP | 2 | 13.00 | - | 6.00 | - | 7.00 | - | - | - | 12 |
| Wendelbo | | S/LO | 2 | 4.75 | - | 14.00 | - | 9.25 | - | - | - | |
| *Dionysia tapetodes* | D | L/UP | 2 | 11.75 | - | 7.75 | - | 4.00 | - | - | - | 12 |
| Bunge | | S/LO | 2 | 4.75 | - | 11.50 | - | 6.75 | - | - | - | |
| *Faramea occidentalis* | D | L/UP | 50 | 13.35 | 1.51 | 9.48 | 1.25 | 3.87 | 0.85 | 0.16 | 0.12 | 13 |
| (L.) A.Rich. | | S/LO | 50 | 6.78 | 1.09 | 12.94 | 1.11 | 6.17 | 1.66 | 0.28 | 0.16 | |
| *Faramea suerrensis* | D | L/UP | 21 | 7.32 | 0.42 | 5.27 | 0.55 | 2.05 | 0.63 | 1.20 | 0.77 | 14 |
| (Donn.Sm.) Donn.Sm. | | S/LO | 11 | 2.96 | 0.27 | 8.48 | 0.76 | 5.52 | 0.65 | 2.31 | 0.59 | |
| *Gaertnera vaginata* | D | L/UP | 25 | 22.12 | 1.57 | 13.69 | 1.38 | 8.43 | 1.55 | 3.12 | 2.11 | 15 |
| Lam. | | S/LO | 25 | 11.68 | 1.33 | 18.41 | 2.01 | 6.78 | 1.92 | 3.02 | 1.55 | |

Table 1 continued.

| **Species** | **PM** | **M/SL** | ***n*** | **Stigma** | **SD** | **Anther** | **SD** | **HER** | **SD** | **MM** | **SD** | **Ref** |
|---|---|---|---|---|---|---|---|---|---|---|---|---|
| *Galianthe peruviana* | D | L/UP | 50 | 3.80 | 0.57 | 2.17 | 0.42 | 1.61 | 2.40 | - | - | 11 |
| (Pers.) E.L.Cabral | | S/LO | 98 | 2.13 | 0.68 | 3.34 | 0.62 | 1.28 | 1.04 | - | - | |
| *Galianthe valerianoides* | D | L/UP | 72 | 5.03 | 0.57 | 3.46 | 0.46 | 1.57 | 0.57 | - | - | 11 |
| (Cham. & Schltdl.) E.L.Cabral | | S/LO | 38 | 2.76 | 0.52 | 4.94 | 0.85 | 2.17 | 0.63 | - | - | |
| *Gelsemium sempervirens* | D | L/UP | 50 | 28.13 | 3.85 | 14.70 | 2.33 | 13.43 | 4.65 | - | - | 16 |
| (L.) J.St.-Hil. | | S/LO | 98 | 15.87 | 3.63 | 23.73 | 3.04 | 8.24 | 4.60 | - | - | |
| *Glandora diffusa* | D | L/UP | 66 | 9.02 | 1.00 | 5.63 | 0.62 | 3.38 | 0.41 | 1.08 | 0.79 | 17 |
| (Lag.) D.C.Thomas | | S/LO | 32 | 5.48 | 0.70 | 8.93 | 0.88 | 3.45 | 0.23 | 0.76 | 0.56 | |
| *Glandora moroccana* | D | L/UP | 51 | 13.74 | 1.34 | 7.80 | 0.48 | 5.94 | 0.87 | 1.68 | 1.23 | 17 |
| (I.M.Johnst.) D.C.Thomas | | S/LO | 49 | 7.30 | 1.03 | 12.49 | 0.99 | 5.19 | 0.17 | 1.02 | 0.70 | |
| *Glandora oleifolia* | D | L/UP | 27 | 13.30 | 1.15 | 7.78 | 0.31 | 5.52 | 0.85 | 1.11 | 0.75 | 17 |
| (Lapeyr.) D.C.Thomas | | S/LO | 37 | 7.20 | 1.46 | 13.25 | 0.69 | 6.05 | 0.84 | 1.39 | 0.79 | |
| *Glandora prostrata* | SHD | L/UP | 52 | 7.92 | 0.89 | 5.97 | 0.71 | 1.95 | 0.24 | 1.67 | 0.98 | 17 |
| (Loisel.) D.C.Thomas | | S/LO | 48 | 4.10 | 0.68 | 6.32 | 0.63 | 2.22 | 0.14 | 1.90 | 0.91 | |
| *Glandora rosmarinifolia* | D | L/UP | 56 | 13.58 | 1.50 | 7.40 | 0.44 | 6.18 | 1.11 | 1.70 | 1.29 | 17 |
| (Ten.) D.C.Thomas | | S/LO | 40 | 7.73 | 1.17 | 12.69 | 1.23 | 4.95 | 0.46 | 1.02 | 0.80 | |
| *Goniolimon italicum* | D | L/UP | 27 | 7.63 | 0.49 | 6.16 | 0.56 | 1.47 | 0.41 | 0.85 | 0.46 | 18 |
| Tammaro, Pignatti & Frizzi | | S/LO | 20 | 6.09 | 0.50 | 6.98 | 0.40 | 0.89 | 0.43 | 0.51 | 0.39 | |
| *Guettarda platypoda* | SHD | L/UP | 95 | 16.55 | 2.40 | 14.36 | 1.89 | - | - | 2.98 | 2.12 | 19 |
| DC. | | S/LO | 96 | 10.68 | 2.12 | 16.57 | 2.79 | - | - | 3.94 | 2.45 | |

Table 1 continued.

| **Species** | **PM** | **M/SL** | ***n*** | **Stigma** | **SD** | **Anther** | **SD** | **HER** | **SD** | **MM** | **SD** | **Ref** |
|---|---|---|---|---|---|---|---|---|---|---|---|---|
| *Guettarda scabra* | D | L/UP | 38 | 16.90 | 3.30 | 14.00 | 2.50 | 2.00 | 2.00 | - | - | 20 |
| (L.) Vent. | | S/LO | 53 | 13.50 | 2.20 | 17.60 | 3.10 | 4.10 | 0.70 | - | - | |
| *Guettarda speciosa* | D | L/UP | 74 | 32.45 | 3.40 | 27.00 | 3.56 | 2.01 | 1.29 | - | - | 21 |
| L. | | S/LO | 42 | 18.77 | 2.60 | 36.21 | 5.26 | 15.40 | 3.69 | - | - | |
| *Houstonia caerulea* | D | L/UP | 380 | 5.95 | 1.96 | 3.16 | 0.92 | 3.01 | 0.21 | 1.47 | 0.93 | 22, 23 |
| L. | | S/LO | 400 | 3.79 | 1.25 | 6.62 | 2.18 | 3.78 | 0.39 | 0.70 | 0.52 | |
| *Houstonia longifolia* | D | L/UP | 47 | 5.53 | 0.46 | 2.21 | 0.20 | 3.30 | 0.38 | 0.89 | 0.58 | 23 |
| Gaertn. | | S/LO | 42 | 2.34 | 0.29 | 4.65 | 0.43 | 2.32 | 0.35 | 0.31 | 0.21 | |
| *Houstonia procumbens* | D | L/UP | 94 | 8.53 | 0.96 | 5.89 | 0.51 | 2.65 | 0.85 | 1.35 | 1.02 | 23 |
| (Walter ex J.F.Gmel.) Standl. | | S/LO | 81 | 5.34 | 0.76 | 9.13 | 0.75 | 3.83 | 0.90 | 1.19 | 1.00 | |
| *Jasminum fruticans* | D | L/UP | 34 | 11.81 | 1.29 | 6.31 | 0.83 | 5.50 | 1.12 | 3.00 | 1.52 | 24 |
| L. | | S/LO | 33 | 6.18 | 1.04 | 8.85 | 0.95 | 2.67 | 1.05 | 1.07 | 0.78 | |
| *Jasminum malabaricum* | SHD | L/UP | 100 | 17.65 | 2.29 | 10.78 | 1.49 | 7.58 | 2.01 | 4.48 | 2.68 | 25 |
| Wight | | S/LO | 100 | 4.85 | 0.80 | 13.36 | 1.93 | 9.17 | 1.70 | 5.94 | 1.68 | |
| *Jasminum odoratissimum* | D | L/UP | 27 | 13.70 | 3.60 | 11.90 | 1.60 | 3.90 | 0.10 | - | - | 26 |
| L. | | S/LO | 27 | 8.60 | 1.10 | 15.60 | 1.10 | 5.00 | 1.80 | - | - | |
| *Kalmiopsis fragrans* | D | L/UP | 41 | 12.75 | 1.32 | 9.45 | 1.36 | 3.39 | 1.57 | 2.22 | 1.34 | 27 |
| Meinke & Kaye | | S/LO | 41 | 6.36 | 0.88 | 14.71 | 1.10 | 8.35 | 1.35 | 3.10 | 1.58 | |
| *Kalmiopsis leachiana* | SHD | L/UP | 16 | 9.17 | 0.52 | 3.86 | 0.43 | 5.31 | 0.23 | 5.19 | 0.78 | 28 |
| (L.F. Hend.) Rehder | | S/LO | 16 | 2.55 | 0.34 | 3.99 | 0.58 | 1.43 | 0.34 | 1.31 | 0.55 | |

Table 1 continued.

| **Species** | **PM** | **M/SL** | ***n*** | **Stigma** | **SD** | **Anther** | **SD** | **HER** | **SD** | **MM** | **SD** | **Ref** |
|---|---|---|---|---|---|---|---|---|---|---|---|---|
| *Linum aretioides* | D | L/UP | 30 | 5.03 | 0.09 | 2.27 | 0.07 | 1.02 | 0.19 | 1.89 | 1.39 | 29 |
| Boiss. | | S/LO | 30 | 3.07 | 0.04 | 6.05 | 0.07 | 2.70 | 1.04 | 1.38 | 0.95 | |
| *Linum campanulatum* | D | L/UP | 23 | 13.44 | 1.18 | 10.04 | 1.26 | 3.40 | 0.86 | 1.22 | 0.94 | 24 |
| L. | | S/LO | 23 | 8.28 | 1.01 | 13.48 | 1.04 | 5.20 | 1.09 | 1.95 | 1.33 | |
| *Linum grandiflorum* | SHD | L/UP | 25 | 6.16 | 0.86 | 8.91 | 1.27 | 2.75 | 0.68 | 3.12 | 1.35 | 30 |
| Desf. | | S/LO | 26 | 4.01 | 0.49 | 9.27 | 1.11 | 5.27 | 0.86 | 4.90 | 1.34 | |
| *Linum suffruticosum* | D | L/UP | 32 | 6.31 | 0.92 | 4.19 | 0.55 | 2.12 | 0.90 | 0.96 | 0.74 | 31 |
| L. | | S/LO | 32 | 3.64 | 0.89 | 6.79 | 0.67 | 3.14 | 0.30 | 0.96 | 0.66 | |
| *Linum tenue* | D | L/UP | 34 | 6.09 | 0.45 | 4.35 | 0.22 | 1.75 | 0.39 | 0.74 | 0.53 | 31 |
| Desf. | | S/LO | 34 | 2.47 | 0.39 | 6.79 | 0.39 | 4.32 | 0.37 | 1.88 | 0.44 | |
| *Linum tenuifolium* | D | L/UP | 67 | 7.61 | 0.93 | 5.87 | 0.72 | 1.74 | 0.56 | 0.97 | 0.73 | 32 |
| L. | | S/LO | 59 | 5.81 | 0.79 | 7.76 | 0.78 | 1.95 | 0.55 | 0.86 | 0.63 | |
| *Lithodora fruticosa* | SHD | L/UP | 52 | 12.22 | 1.15 | 8.97 | 1.16 | 3.25 | 0.17 | 1.69 | 1.21 | 33 |
| (L.) Griseb. | | S/LO | 48 | 5.81 | 1.00 | 11.12 | 1.33 | 5.31 | 0.36 | 3.19 | 1.48 | |
| *Luculia pinceana* | D | L/UP | 66 | 30.75 | 2.03 | 22.74 | 1.46 | 3.98 | 1.38 | 0.99 | 0.74 | 34 |
| Hook. | | S/LO | 60 | 19.80 | 2.71 | 29.75 | 3.64 | 4.40 | 1.16 | 1.15 | 0.81 | |
| *Melochia nodiflora* | D | L/UP | 27 | 9.56 | 0.55 | 4.83 | 0.30 | 4.73 | 0.54 | 1.11 | 0.61 | 35 |
| Sw. | | S/LO | 11 | 5.34 | 0.46 | 8.47 | 0.71 | 3.14 | 0.42 | 0.58 | 0.36 | |
| *Melochia pyramidata* | D | L/UP | 23 | 4.83 | 0.51 | 3.90 | 0.36 | 0.93 | 0.33 | 0.46 | 0.35 | 35 |
| L. | | S/LO | 17 | 3.58 | 0.28 | 4.67 | 0.60 | 1.10 | 0.27 | 0.70 | 0.51 | |

Table 1 continued.

| **Species** | **PM** | **M/SL** | ***n*** | **Stigma** | **SD** | **Anther** | **SD** | **HER** | **SD** | **MM** | **SD** | **Ref** |
|---|---|---|---|---|---|---|---|---|---|---|---|---|
| *Melochia savannarum* | D | L/UP | 46 | 9.28 | 0.79 | 4.76 | 0.92 | 4.52 | 0.54 | 1.07 | 0.82 | 35 |
| Britt. | | S/LO | 54 | 5.48 | 0.66 | 8.49 | 1.24 | 3.01 | 0.65 | 0.93 | 0.69 | |
| *Melochia tomentosa* | D | L/UP | 39 | 7.55 | 1.32 | 4.37 | 0.69 | 3.18 | 1.05 | 1.51 | 1.03 | 35 |
| L. | | S/LO | 23 | 4.18 | 0.58 | 6.51 | 1.05 | 2.33 | 0.58 | 0.62 | 0.46 | |
| *Melochia villosa* | D | L/UP | 16 | 7.31 | 0.49 | 4.16 | 0.36 | 3.09 | 0.31 | 1.02 | 0.55 | 35 |
| (Mill.) Fawc. & Rendle | | S/LO | 24 | 5.18 | 0.47 | 6.27 | 0.67 | 1.17 | 0.15 | 0.93 | 0.48 | |
| *Menyanthes trifoliata* | D | L/UP | 23 | 12.60 | 1.49 | 6.70 | 0.96 | 5.90 | 0.96 | 2.83 | 1.88 | 36 |
| L. | | S/LO | 17 | 8.70 | 1.15 | 10.70 | 1.61 | 2.10 | 0.95 | 1.63 | 1.14 | |
| *Morinda pubescens* | D | L/UP | 21 | 23.11 | 3.19 | 15.05 | 2.32 | 8.06 | 1.14 | 4.18 | 3.24 | This |
| Sm. | | S/LO | 25 | 11.88 | 2.07 | 24.89 | 3.81 | 13.01 | 1.87 | 3.64 | 2.55 | study |
| *Mussaenda decipiens* | D | L/UP | 25 | 24.90 | 1.48 | 15.06 | 1.14 | 9.84 | 0.91 | 4.14 | 2.11 | 37 |
| H.Li | | S/LO | 20 | 15.19 | 1.19 | 20.80 | 1.69 | 5.61 | 1.67 | 1.28 | 0.98 | |
| *Mussaenda divaricata* | D | L/UP | 25 | 24.36 | 0.94 | 17.89 | 0.73 | 6.47 | 0.80 | 1.24 | 0.81 | 37 |
| Hutch. | | S/LO | 25 | 11.21 | 0.66 | 23.47 | 0.75 | 12.26 | 0.51 | 6.69 | 0.96 | |
| *Mussaenda erosa* | D | L/UP | 25 | 23.12 | 0.88 | 13.47 | 0.54 | 9.65 | 0.95 | 3.70 | 1.52 | 37 |
| Champ. ex Benth. | | S/LO | 25 | 9.75 | 1.03 | 19.42 | 1.28 | 9.67 | 0.91 | 3.72 | 1.14 | |
| *Mussaenda hainanensis* | D | L/UP | 16 | 27.84 | 1.48 | 15.10 | 0.78 | 12.74 | 0.98 | 7.54 | 2.18 | 37 |
| Merr. | | S/LO | 19 | 7.16 | 0.54 | 20.30 | 1.69 | 13.14 | 1.49 | 7.94 | 0.92 | |
| *Mussaenda hirsutula* | D | L/UP | 20 | 23.36 | 1.20 | 16.18 | 1.39 | 7.17 | 0.94 | 3.67 | 1.58 | 37 |
| Miq. | | S/LO | 17 | 12.63 | 0.58 | 19.73 | 1.21 | 7.10 | 0.81 | 3.60 | 1.36 | |

Table 1 continued.

| **Species** | **PM** | **M/SL** | ***n*** | **Stigma** | **SD** | **Anther** | **SD** | **HER** | **SD** | **MM** | **SD** | **Ref** |
|---|---|---|---|---|---|---|---|---|---|---|---|---|
| *Mussaenda kwangsiensis* | D | L/UP | 12 | 20.59 | 3.17 | 12.73 | 1.94 | 7.86 | 1.57 | 3.79 | 2.70 | 37 |
| H.L.Li | | S/LO | 26 | 6.42 | 1.19 | 21.17 | 3.54 | 14.74 | 3.22 | 6.31 | 2.19 | |
| *Mussaenda kwangtungensis* | D | L/UP | 25 | 25.86 | 1.24 | 17.95 | 0.67 | 7.91 | 1.17 | 2.87 | 2.17 | 37 |
| H.L.Li | | S/LO | 25 | 6.95 | 0.48 | 23.53 | 2.51 | 16.58 | 2.25 | 11.01 | 0.81 | |
| *Mussaenda lancipetala* | D | L/UP | 40 | 23.72 | 2.42 | 13.75 | 1.32 | 9.97 | 1.70 | 3.27 | 2.46 | 37 |
| X.F.Deng & D.X.Zhang | | S/LO | 38 | 13.94 | 1.58 | 21.13 | 2.10 | 7.19 | 1.41 | 1.66 | 1.20 | |
| *Mussaenda macrophylla* | D | L/UP | 25 | 20.73 | 1.33 | 11.55 | 0.73 | 9.18 | 1.46 | 2.05 | 1.85 | 37 |
| Wall. | | S/LO | 25 | 15.95 | 1.76 | 19.03 | 1.78 | 3.08 | 1.31 | 4.40 | 1.87 | |
| *Mussaenda mollissima* | D | L/UP | 15 | 26.03 | 3.63 | 19.62 | 3.50 | 6.42 | 1.52 | 4.20 | 3.22 | 37 |
| C.Y.Wu ex H.H.Hsue & H.Wu | | S/LO | 14 | 15.89 | 1.73 | 22.59 | 2.04 | 6.70 | 2.14 | 4.53 | 2.76 | |
| *Mussaenda multinervis* | D | L/UP | 25 | 25.51 | 1.87 | 14.44 | 0.89 | 11.07 | 1.80 | 7.59 | 1.93 | 37 |
| C.Y.Wu ex H.H.Hsue & H.Wu | | S/LO | 25 | 16.65 | 0.79 | 17.92 | 0.62 | 1.27 | 0.44 | 2.22 | 1.15 | |
| *Mussaenda parviflora* | D | L/UP | 32 | 12.40 | 1.18 | 6.46 | 1.01 | 6.02 | 0.79 | 4.18 | 1.66 | 38 |
| Miq. | | S/LO | 22 | 5.76 | 0.75 | 8.28 | 0.98 | 2.51 | 1.11 | 1.01 | 0.83 | |
| *Mussaenda pingbianensis* | D | L/UP | 25 | 25.37 | 1.51 | 18.01 | 1.40 | 7.36 | 0.81 | 1.68 | 1.26 | 37 |
| C.Y.Wu ex H.H.Hsue & H.Wu | | S/LO | 25 | 6.60 | 0.31 | 24.70 | 1.36 | 18.10 | 1.13 | 11.40 | 1.40 | |
| *Mussaenda pubescens* | D | L/UP | 25 | 11.20 | 1.34 | 8.14 | 0.66 | 3.06 | 1.17 | 1.61 | 0.78 | 37 |
| Dryand. | | S/LO | 25 | 4.63 | 0.55 | 10.63 | 0.65 | 6.00 | 0.84 | 4.72 | 0.70 | |
| *Mussaenda pubescens* | SHD | L/UP | 26 | 13.71 | 5.25 | 10.70 | 2.65 | 3.95 | 1.12 | - | - | 39 |
| Dryand. | | S/LO | 26 | 4.54 | 2.24 | 14.64 | 1.99 | 9.17 | 6.02 | - | - | |

Table 1 continued.

| **Species** | **PM** | **M/SL** | ***n*** | **Stigma** | **SD** | **Anther** | **SD** | **HER** | **SD** | **MM** | **SD** | **Ref** |
|---|---|---|---|---|---|---|---|---|---|---|---|---|
| *Narcissus assoanus* | SHD | L/UP | 275 | 24.53 | 2.18 | 22.63 | 2.34 | 0.84 | 1.61 | 2.86 | 2.15 | 28 |
| Dufour ex Schult. & Schult.f. | | S/LO | 53 | 13.15 | 2.18 | 23.15 | 2.34 | 4.91 | 1.61 | 5.34 | 3.32 | |
| *Narcissus broussonetii* | SHD | L/UP | 128 | 37.55 | 3.67 | 29.33 | 3.54 | 7.89 | 3.96 | 6.52 | 3.99 | 40 |
| Lag. | | S/LO | 128 | 24.63 | 3.35 | 31.40 | 2.74 | 13.20 | 3.58 | 5.43 | 3.98 | |
| *Narcissus calcicola* | SHD | L/UP | 81 | 17.84 | 1.41 | 15.11 | 1.76 | 1.20 | 1.30 | - | - | 41 |
| Mendonça | | S/LO | 80 | 8.00 | 1.41 | 15.97 | 1.75 | 3.47 | 1.30 | - | - | |
| *Narcissus cuatrecasasii* | SHD | L/UP | 68 | 15.67 | 1.28 | 16.39 | 1.08 | 0.25 | 0.73 | - | - | 41 |
| Fern.Casas, M.Laínz & Ruíz Rejón | | S/LO | 54 | 8.47 | 1.28 | 17.26 | 1.08 | 2.64 | 0.72 | - | - | |
| *Narcissus dubius* | SHD | L/UP | 450 | 15.62 | 1.72 | 15.42 | 1.23 | 0.08 | - | - | - | 28 |
| Gouan | | S/LO | 272 | 9.38 | 2.97 | 15.79 | 2.82 | 2.66 | - | - | - | |
| *Narcissus gaditanus* | SHD | L/UP | 52 | 18.62 | 1.56 | 15.95 | 1.51 | 1.64 | 1.19 | - | - | 41 |
| Boiss. & Reut. | | S/LO | 51 | 8.10 | 1.59 | 16.24 | 1.51 | 3.16 | 1.19 | - | - | |
| *Narcissus papyraceus* | SHD | L/UP | 33 | 18.06 | 1.60 | 20.39 | 1.71 | 0.03 | 1.06 | 3.61 | 2.33 | 41 |
| Ker Gawl. | | S/LO | 30 | 9.61 | 1.60 | 20.47 | 1.70 | 3.24 | 1.06 | 4.25 | 2.40 | |
| *Narcissus rupicola* | SHD | L/UP | 27 | 15.91 | 1.21 | 20.63 | 1.78 | 1.51 | 1.27 | - | - | 41 |
| Dufour | | S/LO | 75 | 9.21 | 1.20 | 20.10 | 1.78 | 5.22 | 1.27 | - | - | |
| *Nivenia argentea* | D | L/UP | 166 | 25.48 | 2.02 | 20.91 | 1.65 | 5.38 | 0.40 | 3.26 | 2.15 | 42 |
| Goldblatt | | S/LO | 120 | 15.41 | 2.18 | 23.14 | 1.90 | 6.92 | 1.17 | 4.63 | 2.66 | |
| *Nivenia binata* | D | L/UP | 39 | 18.05 | 1.76 | 13.73 | 1.62 | 4.29 | 0.97 | - | - | 42 |
| Klatt | | S/LO | 15 | 12.03 | 1.60 | 16.05 | 1.12 | 4.18 | 0.75 | - | - | |

Table 1 continued.

| **Species** | **PM** | **M/SL** | ***n*** | **Stigma** | **SD** | **Anther** | **SD** | **HER** | **SD** | **MM** | **SD** | **Ref** |
|---|---|---|---|---|---|---|---|---|---|---|---|---|
| *Nivenia corymbosa* | D | L/UP | 102 | 19.46 | 2.20 | 12.24 | 1.28 | 7.67 | 1.42 | 2.18 | 1.67 | 42 |
| (Ker Gawl.) Baker | | S/LO | 104 | 10.68 | 1.27 | 19.89 | 1.84 | 8.93 | 1.16 | 1.69 | 1.20 | |
| *Nivenia inaequalis* | D | L/UP | 158 | 41.38 | 2.95 | 35.25 | 4.74 | 6.95 | 2.00 | 5.09 | 3.54 | 42 |
| Goldblatt & J.C.Manning | | S/LO | 138 | 30.90 | 2.80 | 37.28 | 3.84 | 6.75 | 2.94 | 4.34 | 3.19 | |
| *Nymphoides geminata* | D | L/UP | 20 | 10.26 | 0.54 | 5.86 | 0.22 | 2.61 | 0.45 | - | - | 43, 44 |
| (R.Br.) Kuntze | | S/LO | 20 | 5.88 | 0.27 | 8.07 | 0.76 | 2.19 | 0.58 | - | - | |
| *Nymphoides montana* | D | L/UP | 20 | 10.43 | 0.54 | 5.88 | 0.45 | 4.67 | 0.55 | 2.56 | 1.03 | 43, 44 |
| Aston | | S/LO | 20 | 5.08 | 0.40 | 8.31 | 0.58 | 2.92 | 0.80 | 0.52 | 0.37 | |
| *Ophiorrhiza japonica* | D | L/UP | 37 | 13.29 | 0.82 | 8.02 | 0.78 | 5.27 | 0.54 | 1.12 | 0.78 | 45 |
| Blume | | S/LO | 40 | 8.84 | 0.87 | 13.78 | 0.99 | 4.93 | 0.98 | 1.16 | 0.81 | |
| *Oplonia nannophylla* | D | L/UP | 20 | 11.45 | 1.30 | 7.56 | 1.06 | 3.74 | 0.76 | 1.54 | 1.08 | 46 |
| (Urb.) Stearn | | S/LO | 20 | 7.09 | 1.20 | 11.02 | 1.41 | 3.93 | 0.78 | 1.28 | 0.97 | |
| *Palicourea coriacea* | D | L/UP | 106 | 9.21 | 1.49 | 7.81 | 1.39 | 1.82 | 0.66 | 1.58 | 1.16 | 11 |
| (Cham.) K.Schum. | | S/LO | 118 | 8.07 | 1.34 | 9.83 | 1.53 | 1.97 | 0.67 | 1.67 | 1.23 | |
| *Palicourea crocea* | D | L/UP | 47 | 17.63 | 1.73 | 13.02 | 1.24 | 4.61 | 1.13 | - | - | 47 |
| (Sw.) Schult. | | S/LO | 40 | 12.38 | 1.11 | 18.81 | 0.91 | 6.43 | 1.79 | - | - | |
| *Palicourea croceoides* | D | L/UP | 46 | 14.84 | 1.58 | 11.27 | 1.14 | 3.59 | 1.36 | 2.09 | 1.48 | 9 |
| Ham. | | S/LO | 44 | 9.61 | 2.25 | 13.76 | 1.73 | 4.16 | 1.40 | 2.52 | 1.63 | |
| *Palicourea demissa* | D | L/UP | 30 | 17.10 | 1.10 | 13.30 | 0.82 | 0.10 | 0.02 | - | - | 48 |
| Standl. | | S/LO | 30 | 13.40 | 0.77 | 19.60 | 1.26 | 0.20 | 0.03 | - | - | |

Table 1 continued.

| **Species** | **PM** | **M/SL** | ***n*** | **Stigma** | **SD** | **Anther** | **SD** | **HER** | **SD** | **MM** | **SD** | **Ref** |
|---|---|---|---|---|---|---|---|---|---|---|---|---|
| *Palicourea marcgravii* | D | L/UP | 77 | 18.48 | 1.70 | 13.66 | 1.49 | 4.80 | 1.39 | - | - | 11 |
| A.St.-Hil. | | S/LO | 130 | 14.00 | 1.86 | 19.91 | 3.03 | 5.90 | 2.60 | - | - | |
| *Palicourea officinalis* | D | L/UP | 224 | 12.31 | 1.50 | 9.62 | 1.33 | 2.88 | 1.05 | - | - | 11 |
| Mart. | | S/LO | 174 | 8.30 | 1.27 | 11.88 | 1.74 | 3.54 | 1.52 | - | - | |
| *Palicourea padifolia* | D | L/UP | 677 | 15.91 | 1.20 | 11.48 | 1.08 | 4.77 | 0.97 | 1.18 | 0.85 | 49 |
| (Willd. ex Schult.) C.M.Taylor & Lorence | | S/LO | 743 | 8.38 | 1.10 | 15.65 | 1.03 | 6.23 | 1.13 | 4.07 | 1.07 | |
| *Palicourea rigida* | D | L/UP | 33 | 15.43 | 1.76 | 12.15 | 1.30 | 3.28 | 1.54 | 0.20 | 0.14 | 11 |
| Kunth | | S/LO | 33 | 10.78 | 1.74 | 16.29 | 2.63 | 5.51 | 2.83 | 0.16 | 0.15 | |
| *Palicourea tetragona* | D | L/UP | 60 | 53.70 | 3.49 | 43.40 | 2.79 | 10.30 | 2.94 | 3.57 | 2.82 | 3, 50 |
| (Donn.Sm.) C.M.Taylor | | S/LO | 60 | 45.60 | 3.10 | 54.00 | 2.48 | 8.40 | 2.63 | 4.48 | 3.15 | |
| *Pentanisia angustifolia* | D | L/UP | 26 | 21.26 | 2.45 | 16.76 | 2.19 | 5.48 | 1.37 | 3.19 | 2.35 | 51 |
| (Hochst.) Hochst. | | S/LO | 24 | 15.46 | 1.52 | 20.06 | 2.20 | 4.73 | 1.01 | 2.76 | 1.99 | |
| *Pentanisia prunelloides* | D | L/UP | 26 | 19.36 | 1.89 | 14.66 | 1.48 | 4.49 | 1.00 | 2.66 | 1.86 | 51 |
| (Klotzsch) Walp. | | S/LO | 24 | 13.86 | 1.27 | 17.96 | 1.57 | 4.98 | 2.00 | 1.94 | 1.38 | |
| *Persicaria wugongshanensis* | D | L/UP | 20 | 4.33 | 0.19 | 2.23 | 0.06 | 1.99 | 0.12 | - | - | 52 |
| | | S/LO | 20 | 2.18 | 0.07 | 4.73 | 0.08 | 2.52 | 0.09 | - | - | |
| *Piriqueta cistoides subsp. caroliniana* | D | L/UP | 6 | 8.52 | 1.03 | 4.90 | 0.61 | 3.62 | 1.03 | - | - | 53 |
| (Walter) Arbo | | S/LO | 4 | 4.67 | 0.52 | 8.50 | 1.30 | 3.83 | 0.93 | - | - | |
| *Polygonum hastatosagittatum* | D | L/UP | 35 | 3.82 | 1.11 | 1.98 | 0.70 | 1.84 | 0.68 | 0.21 | 0.14 | 54 |
| Makino | | S/LO | 35 | 1.71 | 0.67 | 3.91 | 0.66 | 2.21 | 0.40 | 0.31 | 0.16 | |

Table 1 continued.

| **Species** | **PM** | **M/SL** | ***n*** | **Stigma** | **SD** | **Anther** | **SD** | **HER** | **SD** | **MM** | **SD** | **Ref** |
|---|---|---|---|---|---|---|---|---|---|---|---|---|
| *Primula algida* | D | L/UP | 2 | 4.00 | - | 2.75 | - | 1.25 | - | - | - | 12 |
| Adams | | S/LO | 5 | 4.00 | 1.41 | 7.10 | 2.86 | 3.10 | 1.47 | - | - | |
| *Primula aliciae* | D | L/UP | 2 | 9.50 | - | 3.50 | - | 6.00 | - | - | - | 12 |
| G. Taylor ex W.W. Sm. & H.R. Fletcher | | S/LO | 2 | 3.50 | - | 9.50 | - | 6.00 | - | - | - | |
| *Primula allionii* | D | L/UP | 40 | 8.25 | 1.16 | 3.51 | 0.62 | 5.49 | 0.58 | 1.56 | 1.10 | 55, 56 |
| Loisel. | | S/LO | 39 | 3.17 | 0.56 | 9.92 | 1.46 | 5.85 | 0.82 | 0.71 | 0.55 | |
| *Primula alpicola* | D | L/UP | 17 | 9.76 | 1.42 | 4.76 | 0.69 | 5.00 | 1.90 | - | - | 12 |
| (W.W. Sm.) Stapf | | S/LO | 17 | 3.88 | 0.84 | 11.62 | 1.36 | 7.74 | 1.19 | - | - | |
| *Primula amethystina* | D | L/UP | 2 | 4.00 | - | 2.50 | - | 1.50 | - | - | - | 12 |
| Franch. | | S/LO | 2 | 1.20 | - | 4.50 | - | 3.30 | - | - | - | |
| *Primula aromatica* | D | L/UP | 2 | 11.50 | - | 3.00 | - | 8.50 | - | - | - | 12 |
| W.W. Sm. & Forrest | | S/LO | 2 | 2.00 | - | 9.00 | - | 7.00 | - | - | - | |
| *Primula asarifolia* | D | L/UP | 2 | 13.00 | - | 4.25 | - | 8.75 | - | - | - | 12 |
| H.R. Fletcher | | S/LO | 2 | 4.00 | - | 12.00 | - | 8.00 | - | - | - | |
| *Primula aurantiaca* | D | L/UP | 27 | 10.67 | 1.12 | 6.94 | 0.81 | 3.72 | 0.67 | - | - | 12 |
| W.W. Sm. & Forrest | | S/LO | 27 | 6.98 | 1.15 | 11.94 | 1.14 | 4.96 | 1.22 | - | - | |
| *Primula auriculata* | D | L/UP | 5 | 8.60 | 0.42 | 6.50 | 0.50 | 2.10 | 0.22 | - | - | 12 |
| Lam. | | S/LO | 5 | 5.20 | 0.45 | 10.20 | 0.45 | 5.00 | - | - | - | |
| *Primula barbicalyx* | D | L/UP | 5 | 13.00 | 2.92 | 3.70 | 0.67 | 9.30 | 2.33 | - | - | 12 |
| C.H. Wright | | S/LO | 5 | 3.10 | 1.02 | 13.40 | 4.08 | 10.30 | 3.09 | - | - | |

Table 1 continued.

| **Species** | **PM** | **M/SL** | ***n*** | **Stigma** | **SD** | **Anther** | **SD** | **HER** | **SD** | **MM** | **SD** | **Ref** |
|---|---|---|---|---|---|---|---|---|---|---|---|---|
| *Primula bella* | D | L/UP | 2 | 8.00 | - | 2.00 | - | 6.00 | - | - | - | 12 |
| Franch. | | S/LO | 2 | 2.00 | - | 8.00 | - | 6.00 | - | - | - | |
| *Primula bellidifolia* | D | L/UP | 22 | 7.66 | 1.43 | 4.05 | 1.30 | 3.61 | 0.62 | - | - | 12 |
| King ex Hook. f. | | S/LO | 31 | 3.61 | 0.91 | 7.61 | 1.31 | 4.00 | 1.47 | - | - | |
| *Primula blattariformis* | D | L/UP | 7 | 11.93 | 1.10 | 6.71 | 1.22 | 5.21 | 2.02 | - | - | 12 |
| Franch. | | S/LO | 2 | 4.00 | - | 10.00 | - | 6.00 | - | - | - | |
| *Primula blinii* | D | L/UP | 2 | 9.00 | - | 2.00 | - | 7.00 | - | - | - | 12 |
| H. Lév. | | S/LO | 2 | 2.00 | - | 9.00 | - | 7.00 | - | - | - | |
| *Primula boreiocalliantha* | D | L/UP | 2 | 11.50 | - | 4.50 | - | 7.00 | - | - | - | 12 |
| Balf. f. & Forrest | | S/LO | 2 | 3.00 | - | 12.50 | - | 9.50 | - | - | - | |
| *Primula boveana* | SHD | L/UP | 22 | 18.17 | 0.92 | 16.61 | 0.89 | 1.56 | 0.73 | - | - | 57 |
| Decne. ex Duby | | S/LO | 9 | 13.04 | 0.59 | 17.38 | 0.57 | 4.34 | 0.47 | - | - | |
| *Primula bracteata* | D | L/UP | 2 | 11.75 | - | 10.50 | - | 1.25 | - | - | - | 12 |
| Franch. | | S/LO | 2 | 3.00 | - | 7.50 | - | 4.50 | - | - | - | |
| *Primula bracteosa* | D | L/UP | 2 | 12.50 | - | 6.25 | - | 6.25 | - | - | - | 12 |
| W. G. Craib | | S/LO | 2 | 8.50 | - | 10.00 | - | 1.50 | - | - | - | |
| *Primula calderiana* | D | L/UP | 2 | 14.00 | - | 6.00 | - | 8.00 | - | - | - | 12 |
| Balf. f. & R.E. Cooper | | S/LO | 2 | 6.50 | - | 11.00 | - | 4.50 | - | - | - | |
| *Primula calliantha* | D | L/UP | 2 | 13.00 | - | 6.50 | - | 6.50 | - | - | - | 12 |
| Franch. | | S/LO | 2 | 3.00 | - | 13.00 | - | 10.00 | - | - | - | |

Table 1 continued.

| **Species** | **PM** | **M/SL** | ***n*** | **Stigma** | **SD** | **Anther** | **SD** | **HER** | **SD** | **MM** | **SD** | **Ref** |
|---|---|---|---|---|---|---|---|---|---|---|---|---|
| *Primula capitata* | D | L/UP | 12 | 6.17 | 1.19 | 4.00 | 1.30 | 2.17 | 0.78 | - | - | 12 |
| Hook. | | S/LO | 12 | 3.25 | 1.08 | 8.13 | 1.19 | 4.88 | 0.57 | - | - | |
| *Primula caveana* | D | L/UP | 2 | 11.00 | - | 3.75 | - | 7.25 | - | - | - | 12 |
| W.W. Sm. | | S/LO | 2 | 5.50 | - | 7.50 | - | 2.00 | - | - | - | |
| *Primula cernua* | D | L/UP | 7 | 9.36 | 0.99 | 2.86 | 0.63 | 6.50 | 1.15 | - | - | 12 |
| Franch. | | S/LO | 7 | 3.21 | 0.99 | 9.07 | 1.17 | 5.86 | 1.44 | - | - | |
| *Primula chapaensis* | D | L/UP | 2 | 8.00 | - | 5.00 | - | 3.00 | - | - | - | 12 |
| Gagnep. | | S/LO | 2 | 4.00 | - | 9.50 | - | 5.50 | - | - | - | |
| *Primula chionantha* | D | L/UP | 2 | 12.00 | - | 4.50 | - | 7.50 | - | - | - | 12 |
| Balf. f. & Forrest | | S/LO | 2 | 3.00 | - | 11.00 | - | 8.00 | - | - | - | |
| *Primula chungensis* | D | L/UP | 23 | 11.46 | 1.04 | 7.72 | 0.72 | 3.70 | 0.67 | 0.86 | 0.69 | 12, 58 |
| Balf. f. & Kingdon-Ward | | S/LO | 14 | 6.67 | 0.72 | 11.36 | 1.36 | 4.71 | 0.87 | 2.50 | 1.81 | |
| *Primula cortusoides* | D | L/UP | 10 | 7.55 | 0.64 | 4.70 | 0.35 | 2.85 | 0.53 | - | - | 12 |
| L. | | S/LO | 10 | 4.40 | 0.39 | 9.60 | 0.46 | 5.20 | 0.59 | - | - | |
| *Primula darialica* | D | L/UP | 8 | 6.38 | 0.35 | 4.50 | 0.27 | 1.88 | 0.23 | - | - | 12 |
| Rupr. | | S/LO | 5 | 4.50 | 0.00 | 6.60 | 0.22 | 2.10 | 0.22 | - | - | |
| *Primula deflexa* | D | L/UP | 13 | 10.35 | 1.64 | 3.85 | 0.69 | 6.50 | 2.17 | - | - | 12 |
| Duthie | | S/LO | 13 | 3.31 | 0.63 | 9.88 | 2.08 | 6.58 | 2.26 | - | - | |
| *Primula dryadifolia* | D | L/UP | 2 | 7.75 | - | 4.00 | - | 3.75 | - | - | - | 12 |
| Franch. | | S/LO | 2 | 4.00 | - | 7.75 | - | 3.75 | - | - | - | |

Table 1 continued.

| **Species** | **PM** | **M/SL** | ***n*** | **Stigma** | **SD** | **Anther** | **SD** | **HER** | **SD** | **MM** | **SD** | **Ref** |
|---|---|---|---|---|---|---|---|---|---|---|---|---|
| *Primula efarinosa* | D | L/UP | 5 | 7.60 | 1.02 | 4.40 | 0.96 | 3.20 | 0.27 | - | - | 12 |
| Pax | | S/LO | 5 | 2.60 | 1.47 | 6.70 | 1.04 | 4.10 | 1.39 | - | - | |
| *Primula elliptica* | D | L/UP | 3 | 9.00 | 0.87 | 4.17 | 0.29 | 4.83 | 0.76 | - | - | 12 |
| Royle | | S/LO | 3 | 4.00 | 0.50 | 9.83 | 0.29 | 5.83 | 0.29 | - | - | |
| *Primula erratica* | D | L/UP | 11 | 5.86 | 0.23 | 3.59 | 0.92 | 2.27 | 0.96 | - | - | 12 |
| W.W. Sm. | | S/LO | 5 | 2.90 | 0.82 | 6.20 | 0.27 | 3.30 | 0.67 | - | - | |
| *Primula exscapa* | D | L/UP | 2 | 13.00 | - | 4.00 | - | 9.00 | - | - | - | 12 |
| Hegetschw. | | S/LO | 2 | 3.50 | - | 10.00 | - | 6.50 | - | - | - | |
| *Primula farinosa* | D | L/UP | 7 | 4.57 | 1.13 | 3.43 | 0.67 | 1.29 | 0.49 | - | - | 12 |
| L. | | S/LO | 7 | 2.84 | 1.12 | 5.29 | 1.47 | 2.44 | 1.93 | - | - | |
| *Primula fasciculata* | D | L/UP | 13 | 7.50 | 1.17 | 5.31 | 1.07 | 2.19 | 0.75 | - | - | 12 |
| Balf. f. & Kingdon-Ward | | S/LO | 12 | 4.63 | 0.80 | 7.54 | 1.30 | 2.92 | 0.93 | - | - | |
| *Primula firmipes* | D | L/UP | 25 | 10.28 | 0.72 | 3.96 | 0.75 | 6.32 | 0.92 | - | - | 12 |
| Balf. f. & Forrest | | S/LO | 18 | 3.78 | 1.14 | 10.25 | 1.22 | 6.47 | 0.96 | - | - | |
| *Primula flaccida* | D | L/UP | 17 | 10.38 | 1.24 | 4.18 | 0.92 | 6.21 | 1.13 | - | - | 12 |
| N.P. Balakr. | | S/LO | 17 | 3.97 | 1.01 | 10.88 | 1.10 | 6.91 | 1.09 | - | - | |
| *Primula floribunda* | D | L/UP | 42 | 10.60 | 1.61 | 5.82 | 1.09 | 4.77 | 1.59 | - | - | 12 |
| Wall. | | S/LO | 33 | 5.82 | 1.50 | 12.05 | 2.01 | 6.23 | 2.79 | - | - | |
| *Primula forbesii* | D | L/UP | 2 | 3.00 | - | 1.50 | - | 1.50 | - | - | - | 12 |
| Franch. | | S/LO | 2 | 1.00 | - | 3.00 | - | 2.00 | - | - | - | |

Table 1 continued.

| **Species** | **PM** | **M/SL** | ***n*** | **Stigma** | **SD** | **Anther** | **SD** | **HER** | **SD** | **MM** | **SD** | **Ref** |
|---|---|---|---|---|---|---|---|---|---|---|---|---|
| *Primula forrestii* | D | L/UP | 2 | 9.00 | - | 5.50 | - | 3.50 | - | - | - | 12 |
| Balf. f. | | S/LO | 2 | 2.50 | - | 9.00 | - | 6.50 | - | - | - | |
| *Primula gemmifera* | D | L/UP | 10 | 11.20 | 1.40 | 7.05 | 1.48 | 4.15 | 0.94 | - | - | 12 |
| Batalin | | S/LO | 10 | 6.45 | 1.14 | 12.05 | 1.48 | 5.60 | 1.22 | - | - | |
| *Primula geraniifolia* | D | L/UP | 16 | 11.16 | 1.31 | 4.59 | 1.21 | 6.56 | 2.10 | - | - | 12 |
| Hook. f. | | S/LO | 13 | 4.27 | 1.24 | 11.15 | 2.42 | 6.88 | 2.78 | - | - | |
| *Primula glabra* | D | L/UP | 7 | 3.79 | 0.76 | 2.64 | 0.63 | 1.14 | 0.63 | - | - | 12 |
| Klatt | | S/LO | 7 | 2.29 | 0.39 | 3.93 | 0.79 | 1.64 | 0.48 | - | - | |
| *Primula glomerata* | D | L/UP | 2 | 8.25 | - | 2.00 | - | 6.25 | - | - | - | 12 |
| Pax | | S/LO | 2 | 2.00 | - | 8.25 | - | 6.25 | - | - | - | |
| *Primula heucherifolia* | D | L/UP | 5 | 13.10 | 1.14 | 3.40 | 1.29 | 9.70 | 1.96 | - | - | 12 |
| Franch. | | S/LO | 5 | 3.60 | 0.55 | 12.70 | 2.28 | 9.10 | 1.82 | - | - | |
| *Primula involucrata* | D | L/UP | 26 | 11.10 | 1.41 | 5.96 | 1.23 | 5.13 | 1.20 | - | - | 12 |
| Wall. ex Duby | | S/LO | 26 | 5.85 | 1.13 | 11.71 | 1.01 | 5.87 | 1.06 | - | - | |
| *Primula latisecta* | D | L/UP | 2 | 11.50 | - | 2.00 | - | 9.50 | - | - | - | 12 |
| W.W. Sm. | | S/LO | 2 | 2.50 | - | 8.50 | - | 6.00 | - | - | - | |
| *Primula littledalei* | D | L/UP | 2 | 8.50 | - | 2.50 | - | 6.00 | - | - | - | 12 |
| Balf. f. & Watt | | S/LO | 2 | 2.50 | - | 8.50 | - | 6.00 | - | - | - | |
| *Primula luteola* | D | L/UP | 5 | 13.00 | 0.61 | 8.10 | 0.22 | 4.90 | 0.65 | - | - | 12 |
| Rupr. | | S/LO | 6 | 4.75 | 0.52 | 12.25 | 0.99 | 7.50 | 0.95 | - | - | |

Table 1 continued.

| **Species** | **PM** | **M/SL** | ***n*** | **Stigma** | **SD** | **Anther** | **SD** | **HER** | **SD** | **MM** | **SD** | **Ref** |
|---|---|---|---|---|---|---|---|---|---|---|---|---|
| *Primula malacoides* | D | L/UP | 20 | 5.73 | 1.37 | 3.63 | 1.13 | 2.10 | 0.80 | - | - | 12 |
| Franch. | | S/LO | 20 | 3.70 | 1.40 | 5.68 | 1.09 | 1.98 | 0.97 | - | - | |
| *Primula malvacea* | D | L/UP | 7 | 10.43 | 2.26 | 5.57 | 0.73 | 4.86 | 2.15 | - | - | 12 |
| Franch. | | S/LO | 2 | 7.50 | - | 10.00 | - | 2.50 | - | - | - | |
| *Primula marginata* | D | L/UP | 13 | 9.03 | 1.50 | 2.79 | 0.59 | 5.67 | 1.20 | 1.78 | 1.25 | 56 |
| Curtis | | S/LO | 13 | 3.43 | 0.62 | 9.11 | 1.41 | 5.34 | 1.32 | 0.74 | 0.55 | |
| *Primula maximowiczii* | D | L/UP | 2 | 11.00 | - | 4.50 | - | 6.50 | - | - | - | 12 |
| Regel | | S/LO | 2 | 3.50 | - | 14.50 | - | 11.00 | - | - | - | |
| *Primula membranifolia* | D | L/UP | 5 | 9.30 | 2.80 | 3.70 | 1.15 | 5.60 | 1.71 | - | - | 12 |
| Franch. | | S/LO | 5 | 3.70 | 1.15 | 9.00 | 2.52 | 5.30 | 1.44 | - | - | |
| *Primula merrilliana* | D | L/UP | 20 | 7.10 | 2.99 | 4.45 | 1.43 | 2.58 | 0.52 | 0.47 | 0.38 | 59 |
| Schltr. | | S/LO | 20 | 4.34 | 1.48 | 7.33 | 1.61 | 2.94 | 0.52 | 0.45 | 0.27 | |
| *Primula minor* | D | L/UP | 2 | 10.00 | - | 5.00 | - | 5.00 | - | - | - | 12 |
| Balf. f. & Kingdon-Ward | | S/LO | 2 | 3.75 | - | 10.50 | - | 6.75 | - | - | - | |
| *Primula mistassinica* | D | L/UP | 297 | 3.72 | 0.38 | 1.93 | 0.24 | 1.20 | 0.33 | 0.60 | 0.46 | 60 |
| Michx. | | S/LO | 295 | 1.99 | 0.26 | 3.30 | 0.40 | 0.65 | 0.38 | 0.39 | 0.30 | |
| *Primula modesta* | D | L/UP | 5 | 5.20 | 0.27 | 4.00 | 0.00 | 1.20 | 0.27 | - | - | 12 |
| Bisset & S.Moore | | S/LO | 5 | 3.90 | 0.22 | 6.10 | 0.22 | 2.20 | 0.27 | - | - | |
| *Primula mollis* | D | L/UP | 5 | 10.00 | 1.22 | 5.70 | 0.45 | 4.30 | 0.84 | - | - | 12 |
| Nutt. ex Hook. | | S/LO | 2 | 6.00 | - | 9.00 | - | 3.00 | - | - | - | |

Table 1 continued.

| **Species** | **PM** | **M/SL** | ***n*** | **Stigma** | **SD** | **Anther** | **SD** | **HER** | **SD** | **MM** | **SD** | **Ref** |
|---|---|---|---|---|---|---|---|---|---|---|---|---|
| *Primula moupinensis* | D | L/UP | 2 | 13.50 | - | 6.00 | - | 7.50 | - | - | - | 12 |
| Franch. | | S/LO | 2 | 5.50 | - | 7.75 | - | 2.25 | - | - | - | |
| *Primula muscarioides* | D | L/UP | 17 | 8.26 | 1.80 | 3.94 | 0.79 | 4.32 | 2.24 | - | - | 12 |
| Hemsl. | | S/LO | 15 | 3.40 | 1.61 | 8.63 | 0.77 | 5.23 | 1.55 | - | - | |
| *Primula nivalis* | D | L/UP | 2 | 9.50 | - | 3.50 | - | 6.00 | - | - | - | 12 |
| Pall. | | S/LO | 2 | 2.00 | - | 9.50 | - | 7.50 | - | - | - | |
| *Primula nutans* | D | L/UP | 8 | 9.69 | 1.41 | 6.31 | 1.56 | 3.38 | 1.83 | - | - | 12 |
| Georgi | | S/LO | 8 | 5.94 | 0.98 | 8.63 | 1.75 | 2.69 | 0.92 | - | - | |
| *Primula obconica* | D | L/UP | 10 | 11.50 | 1.85 | 3.85 | 1.06 | 7.19 | 1.65 | - | - | 12 |
| Hance | | S/LO | 9 | 4.05 | 0.96 | 11.61 | 1.96 | 7.17 | 1.71 | - | - | |
| *Primula odontocalyx* | D | L/UP | 2 | 9.50 | - | 4.75 | - | 4.75 | - | - | - | 12 |
| (Franch.) Pax | | S/LO | 2 | 7.50 | - | 8.50 | - | 1.00 | - | - | - | |
| *Primula orbicularis* | D | L/UP | 2 | 8.75 | - | 4.50 | - | 4.25 | - | - | - | 12 |
| Hemsl. | | S/LO | 2 | 4.50 | - | 8.75 | - | 4.25 | - | - | - | |
| *Primula oreodoxa* | D | L/UP | 111 | 9.68 | 0.78 | 5.73 | 0.53 | 3.95 | 0.94 | 0.93 | 0.67 | 61 |
| Franch. | | S/LO | 117 | 5.78 | 0.56 | 9.70 | 0.84 | 3.92 | 0.77 | 0.61 | 0.47 | |
| *Primula ovalifolia* | D | L/UP | 2 | 9.00 | - | 4.50 | - | 4.50 | - | - | - | 12 |
| Franch. | | S/LO | 2 | 4.25 | - | 11.00 | - | 6.75 | - | - | - | |
| *Primula palinuri* | D | L/UP | 30 | 13.30 | 1.16 | 7.50 | 1.08 | 5.80 | 1.69 | - | - | 62 |
| Petagna | | S/LO | 30 | 7.40 | 1.07 | 13.60 | 1.90 | 6.20 | 2.30 | - | - | |

Table 1 continued.

| **Species** | **PM** | **M/SL** | ***n*** | **Stigma** | **SD** | **Anther** | **SD** | **HER** | **SD** | **MM** | **SD** | **Ref** |
|---|---|---|---|---|---|---|---|---|---|---|---|---|
| *Primula partschiana* | D | L/UP | 2 | 11.00 | - | 5.50 | - | 5.50 | - | - | - | 12 |
| Pax | | S/LO | 2 | 5.50 | - | 11.00 | - | 5.50 | - | - | - | |
| *Primula pinnatifida* | D | L/UP | 12 | 8.25 | 0.66 | 3.04 | 0.62 | 5.21 | 0.50 | - | - | 12 |
| Franch. | | S/LO | 7 | 2.43 | 0.35 | 8.29 | 0.81 | 5.86 | 0.63 | - | - | |
| *Primula poissonii* | D | L/UP | 60 | 7.85 | 0.58 | 3.83 | 0.30 | 4.02 | 0.56 | - | - | 63 |
| Franch. | | S/LO | 60 | 3.24 | 0.33 | 8.11 | 0.67 | 4.87 | 0.63 | - | - | |
| *Primula polyneura* | D | L/UP | 33 | 13.53 | 1.47 | 8.24 | 1.23 | 5.29 | 1.08 | - | - | 12 |
| Franch. | | S/LO | 33 | 7.43 | 1.36 | 13.67 | 1.57 | 6.24 | 0.87 | - | - | |
| *Primula primulina* | D | L/UP | 2 | 4.00 | - | 2.00 | - | 2.00 | - | - | - | 12 |
| (Spreng.) H. Hara | | S/LO | 2 | 2.00 | - | 4.00 | - | 2.00 | - | - | - | |
| *Primula prolifera* | D | L/UP | 46 | 9.53 | 1.46 | 4.27 | 0.70 | 5.29 | 1.39 | - | - | 12 |
| Wall. | | S/LO | 37 | 4.12 | 0.45 | 11.31 | 1.88 | 7.19 | 1.88 | - | - | |
| *Primula pulchella* | D | L/UP | 5 | 9.90 | 1.43 | 4.10 | 1.95 | 5.80 | 2.46 | - | - | 12 |
| Franch. | | S/LO | 3 | 3.33 | 2.31 | 10.00 | 2.00 | 6.67 | 3.06 | - | - | |
| *Primula pulverulenta* | D | L/UP | 17 | 13.65 | 1.09 | 9.29 | 1.02 | 4.35 | 0.49 | - | - | 12 |
| Duthie | | S/LO | 22 | 8.32 | 1.28 | 14.59 | 1.27 | 6.27 | 0.53 | - | - | |
| *Primula pumilio* | D | L/UP | 2 | 4.50 | - | 2.25 | - | 2.25 | - | - | - | 12 |
| Maxim. | | S/LO | 2 | 2.25 | - | 4.50 | - | 2.25 | - | - | - | |
| *Primula pycnoloba* | D | L/UP | 2 | 12.00 | - | 7.00 | - | 5.00 | - | - | - | 12 |
| Bureau & Franch. | | S/LO | 2 | 9.50 | - | 18.00 | - | 8.50 | - | - | - | |

Table 1 continued.

| **Species** | **PM** | **M/SL** | ***n*** | **Stigma** | **SD** | **Anther** | **SD** | **HER** | **SD** | **MM** | **SD** | **Ref** |
|---|---|---|---|---|---|---|---|---|---|---|---|---|
| *Primula reidii* | SHD | L/UP | 7 | 9.57 | 1.17 | 6.86 | 1.11 | 2.71 | 0.39 | - | - | 12 |
| Duthie | | S/LO | 2 | 4.00 | - | 8.00 | - | 4.00 | - | - | - | |
| *Primula reticulata* | D | L/UP | 22 | 12.57 | 1.68 | 7.48 | 2.16 | 5.09 | 1.43 | - | - | 12 |
| Wall. | | S/LO | 10 | 3.95 | 1.04 | 11.20 | 1.06 | 7.25 | 1.40 | - | - | |
| *Primula rotundifolia* | D | L/UP | 2 | 11.00 | - | 5.50 | - | 5.50 | - | - | - | 12 |
| Wall. | | S/LO | 2 | 5.50 | - | 11.00 | - | 5.50 | - | - | - | |
| *Primula rugosa* | D | L/UP | 2 | 7.00 | - | 3.50 | - | 3.50 | - | - | - | 12 |
| N.P. Balakr. | | S/LO | 2 | 3.50 | - | 7.00 | - | 3.50 | - | - | - | |
| *Primula rupestris* | D | L/UP | 2 | 9.25 | - | 6.50 | - | 2.75 | - | - | - | 12 |
| Balf. f. & Farrer | | S/LO | 2 | 5.25 | - | 8.50 | - | 3.25 | - | - | - | |
| *Primula saturata* | D | L/UP | 2 | 13.00 | - | 7.00 | - | 6.00 | - | - | - | 12 |
| W.W. Sm. & H.R. Fletcher | | S/LO | 2 | 5.00 | - | 8.50 | - | 3.50 | - | - | - | |
| *Primula secundiflora* | D | L/UP | 60 | 11.31 | 0.76 | 5.07 | 0.44 | 6.23 | 0.72 | - | - | 63 |
| Franch. | | S/LO | 60 | 4.73 | 0.46 | 11.58 | 0.79 | 6.84 | 0.80 | - | - | |
| *Primula serratifolia* | D | L/UP | 23 | 11.17 | 1.00 | 6.09 | 1.14 | 5.08 | 1.16 | - | - | 12 |
| Franch. | | S/LO | 21 | 5.48 | 1.17 | 11.29 | 1.35 | 5.81 | 0.86 | - | - | |
| *Primula sertulum* | D | L/UP | 2 | 7.50 | - | 2.00 | - | 5.50 | - | - | - | 12 |
| Franch. | | S/LO | 2 | 2.00 | - | 7.50 | - | 5.50 | - | - | - | |
| *Primula sinensis* | D | L/UP | 2 | 13.50 | - | 6.75 | - | 6.75 | - | - | - | 12 |
| Sabine ex Lindl. | | S/LO | 2 | 6.75 | - | 13.50 | - | 6.75 | - | - | - | |

Table 1 continued.

| **Species** | **PM** | **M/SL** | ***n*** | **Stigma** | **SD** | **Anther** | **SD** | **HER** | **SD** | **MM** | **SD** | **Ref** |
|---|---|---|---|---|---|---|---|---|---|---|---|---|
| *Primula sinolisteri* | D | L/UP | 5 | 11.20 | 1.64 | 4.50 | 0.71 | 6.70 | 0.97 | - | - | 12 |
| Balf. f. | | S/LO | 5 | 4.20 | 1.10 | 11.00 | 2.74 | 6.80 | 1.64 | - | - | |
| *Primula sinomollis* | D | L/UP | 2 | 5.00 | - | 3.00 | - | 2.00 | - | - | - | 12 |
| Balf. f. & Forrest | | S/LO | 2 | 4.75 | - | 6.50 | - | 1.75 | - | - | - | |
| *Primula sonchifolia* | D | L/UP | 2 | 10.50 | - | 4.75 | - | 5.75 | - | - | - | 12 |
| Franch. | | S/LO | 2 | 6.00 | - | 11.00 | - | 5.00 | - | - | - | |
| *Primula soongii* | D | L/UP | 2 | 13.00 | - | 6.50 | - | 6.50 | - | - | - | 12 |
| F.H. Chen & C.M. Hu | | S/LO | 2 | 7.50 | - | 14.00 | - | 6.50 | - | - | - | |
| *Primula souliei* | D | L/UP | 2 | 6.50 | - | 2.50 | - | 4.00 | - | - | - | 12 |
| Franch. | | S/LO | 2 | 1.00 | - | 6.00 | - | 5.00 | - | - | - | |
| *Primula tenuiloba* | D | L/UP | 2 | 3.50 | - | 2.00 | - | 1.50 | - | - | - | 12 |
| (Watt) Pax | | S/LO | 2 | 2.00 | - | 3.50 | - | 1.50 | - | - | - | |
| *Primula veris* | D | L/UP | 192 | 15.57 | 1.16 | 8.79 | 0.88 | 6.54 | 1.37 | 3.10 | 1.81 | 64, 65 |
| L. | | S/LO | 199 | 8.94 | 0.86 | 14.82 | 1.18 | 7.22 | 1.48 | 1.94 | 1.44 | |
| *Primula vialii* | D | L/UP | 10 | 8.05 | 0.86 | 3.40 | 1.24 | 4.65 | 1.16 | - | - | 12 |
| Delavay ex Franch. | | S/LO | 15 | 3.27 | 1.13 | 8.63 | 0.64 | 5.37 | 1.20 | - | - | |
| *Primula violacea* | D | L/UP | 3 | 8.67 | 0.29 | 4.67 | 0.29 | 4.00 | - | - | - | 12 |
| W.W. Sm. & Kingdon-Ward | | S/LO | 3 | 2.83 | 0.29 | 9.83 | 0.29 | 7.00 | - | - | - | |
| *Primula vulgaris* | D | L/UP | 188 | 15.89 | 1.27 | 9.33 | 1.05 | 6.81 | 1.15 | 1.60 | 1.23 | 64 |
| Huds. | | S/LO | 186 | 8.53 | 0.76 | 15.73 | 1.45 | 5.88 | 1.18 | 2.61 | 1.38 | |

Table 1 continued.

| **Species** | **PM** | **M/SL** | ***n*** | **Stigma** | **SD** | **Anther** | **SD** | **HER** | **SD** | **MM** | **SD** | **Ref** |
|---|---|---|---|---|---|---|---|---|---|---|---|---|
| *Primula walshii* | D | L/UP | 2 | 3.75 | - | 1.75 | - | 2.00 | - | - | - | 12 |
| W. G. Craib | | S/LO | 2 | 1.75 | - | 3.75 | - | 2.00 | - | - | - | |
| *Primula waltonii* | D | L/UP | 24 | 10.15 | 1.16 | 4.35 | 0.83 | 5.79 | 1.34 | - | - | 12 |
| Watt ex Balf. f. | | S/LO | 25 | 4.30 | 0.94 | 11.80 | 1.24 | 7.50 | 0.69 | - | - | |
| *Primula watsonii* | D | L/UP | 5 | 9.20 | 1.30 | 5.30 | 1.15 | 3.90 | 0.65 | - | - | 12 |
| Dunn | | S/LO | 2 | 4.25 | - | 8.50 | - | 4.25 | - | - | - | |
| *Primula wigramiana* | D | L/UP | 7 | 11.64 | 0.63 | 6.07 | 0.67 | 5.57 | 1.02 | - | - | 12 |
| W.W.Sm. | | S/LO | 3 | 5.17 | 0.58 | 10.50 | 1.00 | 5.33 | 0.58 | - | - | |
| *Primula yunnanensis* | D | L/UP | 8 | 8.81 | 0.65 | 3.19 | 0.88 | 5.63 | 1.25 | - | - | 12 |
| Franch. | | S/LO | 8 | 3.00 | 1.00 | 8.75 | 1.00 | 5.94 | 1.29 | - | - | |
| *Psychotria asiatica* | D | L/UP | 11 | 3.30 | 0.22 | 2.70 | 0.32 | 0.52 | 0.26 | 0.37 | 0.26 | 66 |
| L. | | S/LO | 18 | 2.14 | 0.28 | 3.60 | 0.29 | 1.34 | 0.36 | 0.96 | 0.43 | |
| *Psychotria boninensis* | D | L/UP | 20 | 3.70 | 0.40 | 2.40 | 0.30 | 1.39 | 0.30 | 0.54 | 0.41 | 67 |
| Nakai | | S/LO | 21 | 1.90 | 0.30 | 4.20 | 0.50 | 2.20 | 0.43 | 0.48 | 0.32 | |
| *Psychotria capitata* | D | L/UP | 73 | 9.92 | 1.04 | 6.42 | 0.88 | 3.46 | 0.77 | 1.21 | 0.77 | 11 |
| Ruiz & Pav. | | S/LO | 46 | 5.84 | 0.84 | 9.42 | 1.07 | 3.60 | 0.97 | 0.60 | 0.44 | |
| *Psychotria carthagenensis* | D | L/UP | 173 | 4.89 | 0.55 | 3.12 | 0.46 | 1.78 | 0.57 | 0.90 | 0.72 | 11, 68 |
| Jacq. | | S/LO | 12 | 3.10 | 0.52 | 4.67 | 1.09 | 1.57 | 0.90 | 0.60 | 0.45 | |
| *Psychotria cephalophora* | D | L/UP | 10 | 4.90 | 0.48 | 3.30 | 0.32 | 1.63 | 0.19 | 0.47 | 0.36 | 69 |
| Merr. | | S/LO | 10 | 2.70 | 0.32 | 5.00 | 0.38 | 2.32 | 0.10 | 0.66 | 0.42 | |

Table 1 continued.

| **Species** | **PM** | **M/SL** | ***n*** | **Stigma** | **SD** | **Anther** | **SD** | **HER** | **SD** | **MM** | **SD** | **Ref** |
|---|---|---|---|---|---|---|---|---|---|---|---|---|
| *Psychotria colorata* | D | L/UP | 38 | 16.68 | 1.60 | 11.45 | 1.15 | 5.23 | 1.44 | 1.82 | 1.32 | 70 |
| (Willd. ex Schult.) Müll.Arg. | | S/LO | 37 | 11.43 | 1.16 | 16.66 | 1.62 | 5.23 | 1.46 | 1.23 | 1.04 | |
| *Psychotria conjugens* | D | L/UP | 8 | 7.30 | 0.42 | 4.70 | 0.63 | 2.70 | 0.59 | - | - | 71 |
| Müll.Arg. | | S/LO | 16 | 4.20 | 0.42 | 7.02 | 0.64 | 2.86 | 0.68 | - | - | |
| *Psychotria deflexa* | D | L/UP | 40 | 6.11 | 0.75 | 3.90 | 0.47 | 2.21 | 0.55 | 1.03 | 0.72 | 72 |
| DC. | | S/LO | 40 | 3.25 | 0.41 | 6.48 | 0.64 | 3.22 | 0.68 | 0.49 | 0.38 | |
| *Psychotria gracilenta* | D | L/UP | 36 | 5.91 | 0.86 | 4.34 | 0.92 | 1.57 | 0.67 | 1.00 | 0.73 | 70 |
| Müll.Arg. | | S/LO | 40 | 3.08 | 0.94 | 5.48 | 0.80 | 2.40 | 0.78 | 1.52 | 0.98 | |
| *Psychotria hastisepala* | D | L/UP | 15 | 10.80 | 1.47 | 9.99 | 1.47 | 2.18 | 0.87 | - | - | 71 |
| Müll.Arg. | | S/LO | 16 | 8.57 | 0.52 | 12.87 | 1.21 | 4.14 | 0.79 | - | - | |
| *Psychotria hoffmannseggiana* | D | L/UP | 43 | 4.60 | 0.60 | 3.00 | 0.50 | 1.73 | 0.81 | 0.63 | 0.49 | 73 |
| (Willd. ex Schult.) Müll.Arg. | | S/LO | 57 | 2.50 | 0.30 | 4.60 | 0.60 | 1.97 | 0.71 | 0.57 | 0.43 | |
| *Psychotria homalosperma* | D | L/UP | 36 | 24.15 | 2.54 | 17.53 | 2.27 | 6.62 | 1.95 | 3.74 | 2.96 | 74 |
| A.Gray | | S/LO | 39 | 11.78 | 2.21 | 22.49 | 3.53 | 5.56 | 2.58 | 5.77 | 3.00 | |
| *Psychotria jasminoides* | D | L/UP | 26 | 11.70 | 0.53 | 7.60 | 0.52 | 4.05 | 0.36 | 1.42 | 1.05 | 75 |
| Standl. | | S/LO | 23 | 7.80 | 0.30 | 11.40 | 0.61 | 3.48 | 0.94 | 1.07 | 1.12 | |
| *Psychotria leiocarpa* | D | L/UP | 45 | 7.85 | 0.69 | 5.38 | 0.57 | 2.47 | 0.53 | - | - | 76 |
| Cham. & Schltdl. | | S/LO | 45 | 5.38 | 0.41 | 7.79 | 0.59 | 2.40 | 0.59 | - | - | |
| *Psychotria mapourioides* | D | L/UP | 73 | 8.61 | 1.12 | 6.27 | 1.06 | 2.33 | 0.75 | - | - | 11 |
| DC. | | S/LO | 77 | 5.84 | 0.74 | 9.10 | 1.30 | 3.26 | 1.01 | - | - | |

Table 1 continued.

| **Species** | **PM** | **M/SL** | ***n*** | **Stigma** | **SD** | **Anther** | **SD** | **HER** | **SD** | **MM** | **SD** | **Ref** |
|---|---|---|---|---|---|---|---|---|---|---|---|---|
| *Psychotria nervosa* | D | L/UP | 30 | 55.40 | 0.88 | 29.40 | 0.66 | 26.03 | 0.92 | 3.60 | 1.87 | 77 |
| Sw. | | S/LO | 30 | 30.80 | 0.93 | 49.30 | 0.99 | 21.23 | 2.25 | 1.73 | 1.24 | |
| *Psychotria poeppigiana* | D | L/UP | 40 | 15.87 | 1.84 | 11.54 | 1.09 | 8.14 | 1.07 | 2.55 | 1.70 | 14, 78 |
| Müll.Arg. | | S/LO | 40 | 7.80 | 1.22 | 15.94 | 1.92 | 7.15 | 0.74 | 1.72 | 1.28 | |
| *Psychotria racemosa* | D | L/UP | 3 | 5.16 | 0.74 | 3.64 | 0.51 | 1.51 | - | - | - | 11 |
| Rich. | | S/LO | 3 | 3.01 | 0.14 | 6.18 | 0.56 | 3.17 | - | - | - | |
| *Psychotria serpens* | D | L/UP | 19 | 4.20 | 0.51 | 2.50 | 0.24 | 1.76 | 0.32 | 0.56 | 0.40 | 79 |
| L. | | S/LO | 24 | 2.20 | 0.22 | 4.50 | 0.38 | 2.30 | 0.15 | 0.35 | 0.23 | |
| *Psychotria suerrensis* | D | L/UP | 14 | 17.11 | 2.05 | 10.01 | 1.07 | 7.11 | 1.33 | 2.28 | 1.90 | 14 |
| Donn.Sm. | | S/LO | 15 | 9.05 | 0.79 | 15.27 | 1.28 | 6.22 | 1.05 | 1.32 | 0.91 | |
| *Psychotria suterella* | D | L/UP | 46 | 16.98 | 0.25 | 12.83 | 0.17 | 5.39 | 0.18 | 1.66 | 1.27 | 80 |
| Müll.Arg. | | S/LO | 30 | 12.05 | 0.18 | 17.43 | 0.30 | 4.14 | 0.15 | 1.43 | 0.99 | |
| *Psychotria trichophora* | D | L/UP | 21 | 12.65 | 1.02 | 9.44 | 0.86 | 3.20 | 0.82 | 1.50 | 1.05 | 73 |
| Müll.Arg. | | S/LO | 22 | 7.79 | 1.16 | 11.84 | 2.04 | 4.05 | 1.53 | 1.35 | 0.87 | |
| *Pulmonaria angustifolia* | D | L/UP | 10 | 8.82 | 1.11 | 4.58 | 0.39 | 4.24 | 1.07 | 0.96 | 0.70 | 81 |
| L. | | S/LO | 10 | 3.74 | 0.73 | 8.91 | 0.57 | 5.17 | 0.92 | 1.00 | 0.58 | |
| *Pulmonaria collina* | D | L/UP | 10 | 9.71 | 1.04 | 4.71 | 0.48 | 5.00 | 1.23 | 1.08 | 0.77 | 81 |
| W.Sauer | | S/LO | 11 | 5.20 | 0.44 | 9.94 | 0.89 | 4.74 | 1.10 | 0.65 | 0.45 | |
| *Pulmonaria longifolia* | D | L/UP | 28 | 11.74 | 1.31 | 5.86 | 0.83 | 5.88 | 1.19 | 1.33 | 0.93 | 24 |
| (Bastard) Boreau | | S/LO | 20 | 5.67 | 0.62 | 11.62 | 1.02 | 5.95 | 0.98 | 0.82 | 0.62 | |

Table 1 continued.

| **Species** | **PM** | **M/SL** | ***n*** | **Stigma** | **SD** | **Anther** | **SD** | **HER** | **SD** | **MM** | **SD** | **Ref** |
|---|---|---|---|---|---|---|---|---|---|---|---|---|
| *Pulmonaria mollis* | D | L/UP | 10 | 10.76 | 1.40 | 5.31 | 0.32 | 5.45 | 1.35 | 1.41 | 1.08 | 81 |
| Wulfen ex Hornem. | | S/LO | 10 | 5.07 | 1.04 | 10.32 | 1.15 | 5.25 | 1.12 | 0.84 | 0.64 | |
| *Pulmonaria obscura* | D | L/UP | 30 | 10.50 | 1.10 | 6.20 | 0.55 | 4.30 | 1.10 | 0.91 | 0.81 | 82 |
| Dumort. | | S/LO | 30 | 6.00 | 0.55 | 10.90 | 0.55 | 5.00 | 0.55 | 0.72 | 0.54 | |
| *Pulmonaria officinalis* | D | L/UP | 102 | 10.42 | 1.03 | 5.33 | 0.50 | 5.09 | 0.97 | 1.07 | 0.81 | 83 |
| L. | | S/LO | 74 | 5.92 | 0.65 | 10.55 | 0.86 | 4.62 | 0.87 | 0.82 | 0.59 | |
| *Pulmonaria saccharata* | D | L/UP | 10 | 12.64 | 1.07 | 5.71 | 0.75 | 6.94 | 1.14 | 1.36 | 0.96 | 81 |
| Mill. | | S/LO | 10 | 5.76 | 1.17 | 11.90 | 1.14 | 6.14 | 1.35 | 1.07 | 0.77 | |
| *Rudgea sessilis* | D | L/UP | 26 | 7.80 | 1.11 | 5.55 | 0.50 | 2.34 | 1.28 | - | - | 71 |
| (Vell.) Müll.Arg. | | S/LO | 25 | 5.05 | 0.90 | 7.85 | 0.47 | 2.92 | 1.16 | - | - | |
| *Salvia brandegeei* | D | L/UP | 50 | 11.01 | 0.71 | 6.85 | 0.64 | 3.87 | 0.52 | 0.93 | 0.73 | 28 |
| Munz | | S/LO | 50 | 8.00 | 0.71 | 11.27 | 0.78 | 3.52 | 0.57 | 1.08 | 0.69 | |
| *Schizomussaenda henryi* | D | L/UP | 25 | 30.07 | 2.02 | 23.17 | 1.79 | 6.90 | 1.31 | 4.19 | 2.39 | 37 |
| (Hutch.) X.F.Deng & D.X.Zhang | | S/LO | 25 | 23.14 | 1.92 | 25.95 | 1.57 | 2.81 | 1.01 | 1.99 | 1.63 | |
| *Solanum torvum* | SHD | L/UP | 13 | 10.08 | 0.74 | 8.93 | 0.65 | 1.15 | 0.69 | 1.59 | 0.74 | 84 |
| Sw. | | S/LO | 17 | 3.40 | 1.33 | 8.53 | 0.39 | 5.10 | 1.43 | 5.52 | 1.44 | |
| *Solanum wrightii* | SHD | L/UP | 14 | 23.86 | 1.17 | 18.73 | 1.17 | 5.13 | 1.44 | 5.33 | 1.42 | 84 |
| Benth. | | S/LO | 14 | 7.64 | 1.75 | 18.52 | 0.80 | 10.88 | 2.23 | 11.09 | 2.11 | |
| *Turnera scabra* | D | L/UP | 30 | 11.80 | 1.10 | 8.40 | 0.80 | 1.30 | 1.00 | 1.74 | 1.12 | 85 |
| Millsp. | | S/LO | 30 | 5.80 | 0.80 | 13.30 | 1.00 | 4.20 | 0.90 | 2.63 | 1.07 | |

Table 1 continued.

| **Species** | **PM** | **M/SL** | ***n*** | **Stigma** | **SD** | **Anther** | **SD** | **HER** | **SD** | **MM** | **SD** | **Ref** |
|---|---|---|---|---|---|---|---|---|---|---|---|---|
| *Turnera subulata* | D | L/UP | 30 | 11.80 | 0.90 | 6.50 | 0.60 | 2.40 | 0.90 | 0.95 | 0.73 | 85 |
| Sm. | | S/LO | 30 | 5.70 | 0.70 | 12.00 | 0.90 | 2.90 | 0.60 | 0.97 | 0.71 | |
| *Turnera ulmifolia* | D | L/UP | 30 | 11.30 | 1.00 | 7.70 | 0.80 | 1.70 | 0.70 | 1.61 | 1.07 | 85 |
| L. | | S/LO | 30 | 5.90 | 0.70 | 12.70 | 0.90 | 3.70 | 0.90 | 1.67 | 0.90 | |
| *Vismia guianensis* | D | L/UP | 10 | 0.70 | 0.03 | 0.41 | 0.03 | 0.29 | 0.05 | 0.03 | 0.02 | 86 |
| (Aubl.) Pers. | | S/LO | 10 | 0.40 | 0.03 | 0.70 | 0.03 | 0.29 | 0.03 | 0.03 | 0.02 | |
| *Waltheria ovata* | D | L/UP | 11 | 7.48 | 0.53 | 5.00 | 0.00 | 4.41 | 0.55 | - | - | 87 |
| Cav. | | S/LO | 10 | 4.28 | 0.26 | 5.00 | 0.00 | 2.32 | 0.43 | - | - | |
