## Supplementary Fig. S1 for "Morph-specific patterns of sex organ positions in species with style length polymorphism"

**Supplementary information**

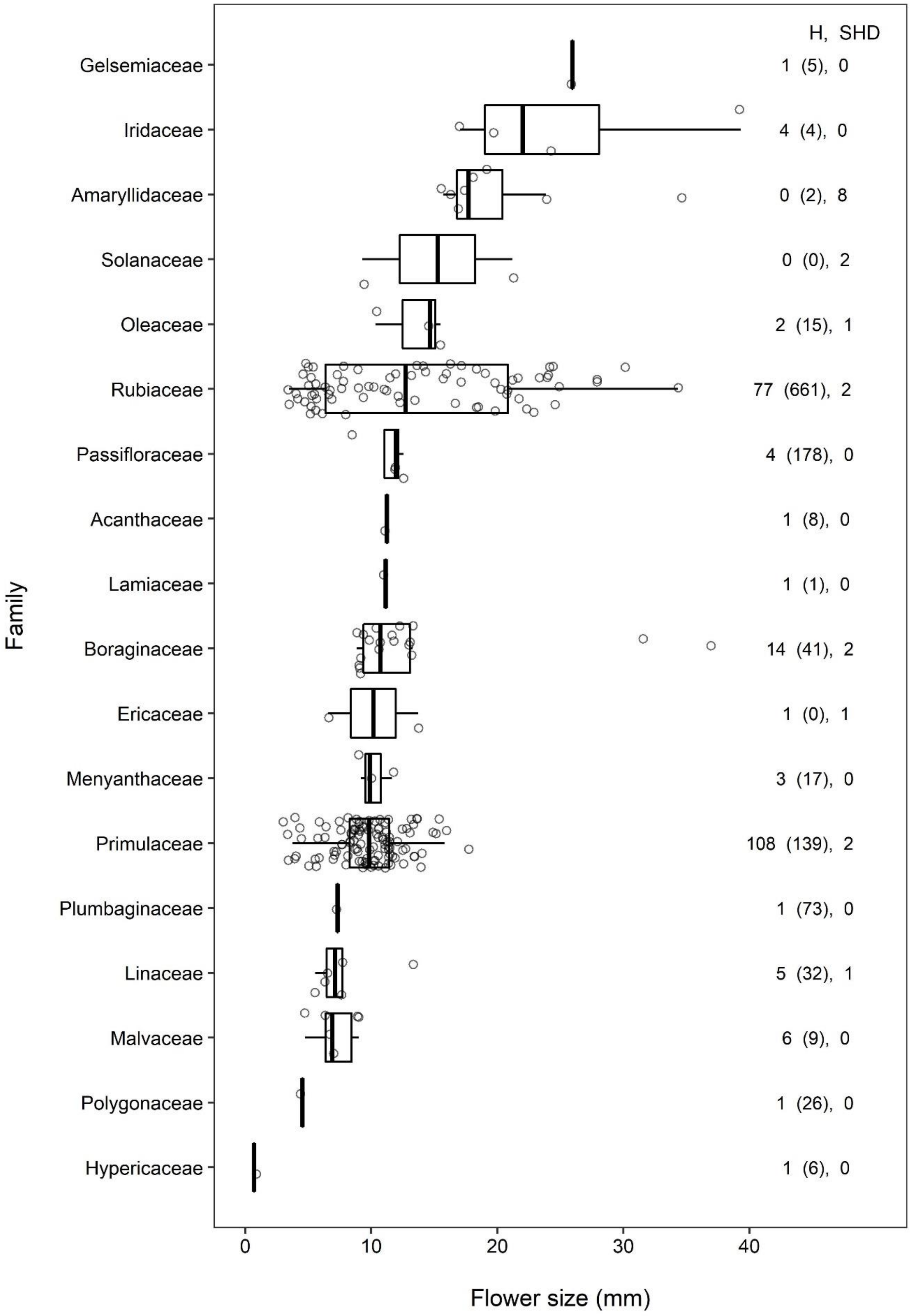

**Figure S1.** The taxonomic distribution of the 230 distylous species and 18 species with stigma-height dimorphism of the current dataset across different families. Values represent the number of distylous species in that family in the current dataset followed by the reported number of heterostylous species in that family (Naiki, 2012), and the number of species with stigma-height dimorphism in the current dataset after a comma. The points represent the mean of the upper level sex organs as a proxy for flower size. The families have been arranged in an ascending order of the median of the proxy for flower size. One species in the family Rubiaceae (*Mussaenda pubescens* Dryand.) has been reported to have heterostyly (distyly), and stigma-height dimorphism in different studies, and hence have been added to both categories in the graph. Three flower sizes above 44 mm have been removed from the family Rubiaceae to help in better visualization of the data. The bold line in the boxplots represents the median, the ends of the box represent the first and the third quartiles and the whiskers represent 1.5 times the interquartile range.
